# Shifts in mutation bias promote mutators by altering the distribution of fitness effects

**DOI:** 10.1101/2022.09.27.509708

**Authors:** Marwa Tuffaha, Saranya Varakunan, David Castellano, Ryan N. Gutenkunst, Lindi M. Wahl

## Abstract

Recent experimental evidence demonstrates that shifts in mutational biases, for example increases in transversion frequency, can change the distribution of fitness effects of mutations (DFE). In particular, reducing or reversing a prevailing bias can increase the probability that a *de novo* mutation is beneficial. It has also been shown that mutator bacteria are more likely to emerge if the beneficial mutations they generate have a larger effect size than observed in the wildtype. Here, we connect these two results, demonstrating that mutator strains that reduce or reverse a prevailing bias have a positively-shifted DFE, which in turn can dramatically increase their emergence probability. Since changes in mutation rate and bias are often coupled through the gain and loss of DNA repair enzymes, our results predict that the invasion of mutator strains will be facilitated by shifts in mutation bias that offer improved access to previously under-sampled beneficial mutations.

## Introduction

*De novo* mutation is foundational to both genetic diversity and adaptive innovation; mutation rates vary both among taxa (Lynch et al., 2016) and across the genome (Hodgkinson and Eyre-Walker, 2011; Martincorena and Luscombe, 2013; Monroe et al., 2022), and can respond rapidly to selective pressure (Wei et al., 2022).

Mutations, however, occur in many forms; single nucleotide substitutions are frequently classified as transitions, transversions, specific base-to-base substitutions or substitutions in *n*-mer contexts, while insertions, deletions, or larger genome rearrangements likewise generate substantial genetic variation (Hodgkinson and Eyre-Walker, 2011). Mutation accumulation studies have allowed for detailed analyses of the rates and contexts of specific types of mutations, the “mutation spectrum” (Katju and Bergthorsson, 2019).

These data have brought to light the fact that the mutation spectrum is biased, in other words, certain classes of mutations occur more frequently than others (Foster et al., 2015; Katju and Bergthorsson, 2019). For example, the frequency of transitions can range from less than 5% to over 95% in bacterial strains that express different DNA repair genes (Foster et al., 2015; Lee et al., 2012), and substantial changes in the mutation spectrum have also been detected in human populations (Harris and Pritchard, 2017). A wealth of recent work has demonstrated that this underlying mutation bias influences and is ultimately echoed in the spectrum of adaptive substitutions (Soares et al., 2021; Stoltzfus and McCandlish, 2017; Stoltzfus and Yampolsky, 2009), even at high mutation rates (Gomez et al., 2020), as demonstrated for fitness landscapes in transcription factor binding sites (Cano and Payne, 2020), for antibiotic resistance (Payne et al., 2019), for convergent mutations in protein sequences (Storz et al., 2019), and for thousands of amino acid changes observed in natural and experimental microbial populations (Cano et al., 2022). As formally derived elsewhere in this special issue (Gitschlag et al., 2023), in the strong-selection-weak-mutation regime fixed substitutions are predicted to be enriched for mutations that occur at high rates. Taken together, this body of work demonstrates that in multiple evolutionary contexts, the spectrum of likely mutations is strongly reflected in the spectrum of adaptive substitutions.

What are the implications of this result? Taking transitions and transversions as an example, the result above implies that beneficial transitions will be more likely to occur and fix, during adaptation, in a population with transition-biased mutations. If there is no *a priori* reason to expect that beneficial mutations are more likely to be transitions or transversions (Stoltzfus and Norris, 2015), this implies that beneficial transversions will be comparatively undersampled as adaptation proceeds. Thus, after a period of adaptation with a transition bias, reducing the bias to sample more transversions, or even reversing it to oversample transversions, can offer access to previously unsampled beneficial mutations. This effect has been recently demonstrated in bacteria, in which shifts in the mutation bias increased the fraction of new mutations that were beneficial (Sane et al., 2021). Simulations of adaptive walks likewise demonstrated a robust effect in which, after a period of adaptation, any reduction or reversal in the existing mutation bias altered the distribution of fitness effects of mutations (DFE), increasing the fraction of beneficial mutations (Sane et al., 2021). This idea, that changing the mutation spectrum could alter (MacLean et al., 2010) or in fact increase the beneficial fraction of the DFE has been previously suggested (Maharjan and Ferenci, 2017), and changes in the DFE have been demonstrated for bacterial mutators evolving antibiotic resistance (Couce et al., 2013).

Mutators, microbial strains that increase the mutation rate by one or several orders of magnitude, are frequently observed in experimental (Miyake, 1960; Shaver et al., 2002; Sniegowski et al., 1997; Treffers et al., 1954; Wei et al., 2022; Wielgoss et al., 2012), natural (Giraud et al., 2001; Jiang et al., 2021; LeClerc et al., 1996; Matic et al., 1997; Oliver et al., 2000; Richardson et al., 2002) and clinical (Couce et al., 2016; Raghavan et al., 2019; Ridderberg et al., 2020) populations; the mutator phenotype is also prevalent in human tumors, which likewise reproduce asexually (Fox et al., 2013). This phenomenon is understood to occur in asexual evolution because a mutation that increases the mutation rate (typically the loss of function of a mismatch repair enzyme (Denamur and Matic, 2006)) hitchhikes with the *de novo* beneficial mutation produced in the mutator strain (Maynard Smith and Haigh, 1974; Sniegowski et al., 1997). The dynamics of mutation rate modifiers have been well-studied both analytically and in simulation; mutators are disfavoured in the presence of genetic exchange (Johnson, 1999; Tenaillon et al., 2000), but favoured in fluctuating environments (Leigh, 1970; Travis and Travis, 2002), or when multiple mutations are required for adaptation either simultaneously (Taddei et al., 1997; Tenaillon et al., 1999) or in sequence (Tanaka et al., 2003). On smooth fitness landscapes, the complex interplay among genetic drift, deleterious load and competition between wildtype and mutator strains is well-understood analytically (Kessler and Levine, 1998; Wylie et al., 2009).

Couce et al. (2013) studied two mutator strains of *E. coli*, each of which increased specific classes of transversions. These changes in mutation spectrum resulted in changes to the DFE for both strains relative to the wildtype. To investigate whether these changes in spectrum could influence mutator emergence, populations were simulated in which the fraction of beneficial mutations was held constant, and every beneficial mutation had fixed effect size *s* in the wildtype and effect size *σs* in mutators. Results predicted that if the effect size of the beneficial mutation in a mutator strain was increased relative to the wildtype, even by a modest amount, the emergence of mutator strains was substantially increased (Couce et al., 2013).

Here, we use theory and simulations to explicitly investigate how a shift in bias affects the DFE, changing both the fraction of mutations that are beneficial and the mean selective effect of beneficial mutations. Using full population simulations, we demonstrate robust benefits of reducing or reversing the mutation bias after a period of adaptation. We investigate the fate of mutations that change the bias, change the mutation rate, or change both the bias and mutation rate. We demonstrate that mutations that reduce or reverse the bias and increase the mutation rate are most likely to emerge, and that this effect is non-linear; a bias shift can dramatically improve the chances of mutator fixation, through the resulting changes in the DFE. Since many loss-of-function mutations affect both mutation rate and bias, our results suggest that shifts in mutation bias may powerfully facilitate the invasion of mutator strains.

## Theory

### The Distribution of Fitness Effects

Suppose an individual with fitness *W* has an offspring that carries a mutation. If the offspring has fitness *W^′^*, the fitness effect of the mutation is defined as *s* = *W^′^*/*W −* 1, where *s* can be negative, zero, or positive. The distribution of fitness effects (DFE) can then be defined as the distribution of such values of *s* from all possible mutations, for a single ancestor genome. This definition, however, assumes that the organism has equal access to all mutations, whereas in reality certain classes of mutations are typically over- or under-represented. It’s therefore important to draw the distinction between the DFE of all possible mutations, DFE_all_, which is often of theoretical interest, and the mutation-weighted (i.e., bias-weighted) DFE, DFE*_β_*. The latter is computed by weighting each entry in DFE_all_ by the rate at which it is expected to occur. We emphasize that DFE*_β_* is the DFE that is accessible experimentally through mutation accumulation experiments, and, critically, accessed by the organism during evolution. In the sections to follow, we will use *f* to denote the fraction of mutations in a DFE that are beneficial, adding subscripts to denote particular cases such as *f*_all_ or *f_β_*. For a list of symbols and their definitions, see Table 1.

**Table 1:** Symbols used in the theoretical analysis.

| Symbol | Definition |
| --- | --- |
| $M$ | Number of possible mutations |
| $B$ | Number of possible beneficial mutations |
| $n$ | Number of fixed mutations after a period of adaptive evolution |
| $\alpha$ | Fraction of mutations of type 1 |
| $\beta$ | Probability that a mutation that occurs is of type 1 (i.e. bias) |
| $\text{DFE}_{\text{all}}$ | Distribution of fitness effects of all possible mutations |
| $\text{DFE}_{\beta}$ | Bias-weighted distribution of fitness effects |
| $f_{\text{all}}$ | Fraction of mutations that are beneficial (“beneficial fraction”) in $\text{DFE}_{\text{all}}$ |
| $f_{\beta}$ | Beneficial fraction in $\text{DFE}_{\beta}$ |
| $f_i$ | Fraction of mutations of type $i$ that are beneficial, $i \in \{1, 2\}$ |
| $f_a$ | Beneficial fraction in the ancestor |
| $\mu$ | Mutation rate per genome per generation |
| $F$ | Mutation rate multiplier in mutator strain |
| $p_{\mu}$ | Fraction of population with higher mutation rate |
| $p_{\text{bs}}$ | Fraction of population with bias shift |
| $p_{\mu, \text{bs}}$ | Fraction of population with higher mutation rate and bias shift |
| $f_{\text{wt}}$ | Beneficial fraction in the wildtype |
| $f_{\text{bs}}$ | Beneficial fraction in the bias-shifted strain |
| $G$ | Increase in beneficial fraction due to the bias shift |
| $\pi_{\text{wt}}$ | Fixation probability of a beneficial mutation in the wildtype |
| $\pi_{\text{bs}}$ | Fixation probability of a beneficial mutation in the bias-shifted strain |
| $H$ | Increase in fixation probability of a beneficial mutation due to the bias shift |
| $d$ | Deleterious fraction in $\text{DFE}_{\beta}$ |
| $s_b$ | Selective advantage of new beneficial mutation |
| $\delta$ | Expected increase in beneficial effect due to a bias shift |
| $\epsilon$ | Expected reduction in deleterious effect due to a bias shift |

A second definition that requires clarification is the concept of unbiased mutation. In each individual, a particular locus in the genome exists in a particular state (for example, as a particular nucleotide, but the argument extends to amino acids or more general “alleles”). A number of other alternate states are possible at that locus. The mutation process is unbiased if each of these alternate states is accessed by mutation with equal probability. Thus unbiased mutation does not depend on the underlying biochemistry of mutation or neutrality of effect; it depends only on the possible alternative states. As an example, a transition fraction of 1/3 is unbiased, because for any nucleotide, 1/3 of the alternate states are reached via transitions. We define mutation classes as over- or under-sampled relative to this unbiased expectation. (We also note that for some classes of mutations, such as GC*→*AT versus AT*→*GC, the unbiased genome-wide mutation rate will depend on genome content; we will not treat these cases here.)

This definition of unbiased mutation also allows us to define bias reinforcements, reductions and reversals, for cases when the mutation bias changes over time. A shift from transition-biased mutation to transversion-biased mutation is a bias reversal. A shift from transition-biased mutation to more extreme transition-biased mutation is a reinforcement. A shift from transition-biased mutation to a less extreme, but still transition-biased mutation process is a bias reduction. In the Introduction, we provided a verbal argument that a reduction or reversal in an existing (historically prevailing) mutation bias increases the beneficial fraction of the DFE*_β_*. Although the logic is straightforward, in the remainder of this section we demonstrate this effect mathematically, which allows us to carefully define the conditions and assumptions under which this expectation holds.

Consider an evolving population in which each individual has *M* possible mutations, in two distinct classes (e.g. transitions and transversions). (The approach below can be generalized to any number of mutation classes but we use two for clarity.) Let *α* be the fraction of possible mutations of type 1, such that there are *αM* mutations in the first class and (1 *− α*)*M* in the second. In the case of transitions as class one and transversions as class two, the value of *α* is 1/3.

Consider a particular “ancestor” genotype. Let *B* denote the total number of beneficial mutations, out of the *M* possible mutations for the ancestor. Thus the fraction of beneficial mutations in the ancestor, defined as *f*_a_, is given by *B*/*M*. Now suppose that we have no *a priori* reason to assume that mutations of one type or another are more likely to be beneficial, nor do they have different beneficial effect sizes. Under the assumption that the beneficial DFEs are the same in the two classes of mutations, we expect *αB* potential beneficial mutations in the first class, and (1 *− α*)*B* in the second, as shown in the first two rows of Table 2.

**Table 2:** Table of values for numbers of mutations and beneficial fractions in the ancestor and in an evolved strain after n fixation events. See text for details.

| number of mutations | total | class 1 (e.g. Ti) | class 2 (e.g. Tv) |
| --- | --- | --- | --- |
| all possible | $M$ | $\alpha M$ | $(1 - \alpha)M$ |
| beneficial, available to ancestor | $B$ | $\alpha B$ | $(1 - \alpha)B$ |
| beneficial, fixed in evolved | $n$ | $\beta n$ | $(1 - \beta)n$ |
| **beneficial fractions** |  |  |  |
| ancestral ($f_a$) | $f_a = \frac{B}{M}$ | $\frac{\alpha B}{\alpha M} = f_a$ | $\frac{(1-\alpha)B}{(1-\alpha)M} = f_a$ |
| evolved, after $n$ fixations | $\frac{B-n}{M} = f_a - \frac{n}{M}$ | $\frac{\alpha B - \beta n}{\alpha M} = f_a - \left(\frac{\beta}{\alpha}\right) \frac{n}{M}$ | $f_a - \frac{(1-\beta)}{(1-\alpha)} \frac{n}{M}$ |

Now assume that after a period of adaptive evolution, *n* beneficial mutations have fixed. For clarity, we will assume that mutations fix sequentially without competition, i.e. the strong-selection weak-mutation regime, but this will be relaxed in the simulations to follow. If there is no mutation bias, beneficial mutations in each class will be equally likely to be sampled and reach fixation, so we expect *αn* fixations in the first mutation class and (1 *− α*)*n* in the second. In contrast, suppose mutation bias has led to a different sampling ratio of mutations. While *α* is the fraction of all possible mutations that are in the first class, let *β* be the fraction of mutations that occur that are in the first class. We would then expect *βn* fixations in the first class, and (1 *− β*)*n* in the second.

Without loss of generality, assume *α < β*, such that class one is the over-sampled class. As a realistic example, we consider a transition-biased organism, such that *β >* 1/3 of fixed mutations are transitions. This implies that mutation bias has led to the fixation of more beneficial mutations of type 1, and fewer of type 2, than expected under the theoretically-defined unbiased mutation process (see Cano et al. (2022) for recent empirical examples).

At this point we must impose further assumptions regarding the fitness landscape. In reality, due to epistasis, each beneficial substitution may change the remaining number of beneficial mutations, both in its own class and in others. Even in the absence of epistasis, a given nucleotide could have two possible beneficial mutations, for example both possible transversions could be beneficial, and after one transversion has fixed, the remaining transversion (now a transition) may or may not provide a further benefit. These complexities are captured in the simulation studies that follow. For analytical tractability, however, we will assume a smooth fitness landscape, and that beneficial mutations are rare. In particular, each locus has at most a single beneficial mutation available.

In this simplified fitness landscape, it is straightforward to compute the fraction of remaining mutations in each class that are beneficial, *f*_1_ and *f*_2_, by simply subtracting the number of beneficial substitutions that have occurred. As shown in the last row of Table 2, we find:

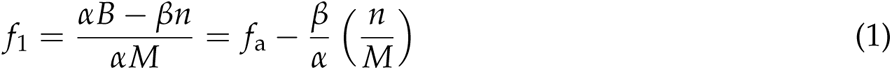

And similarly

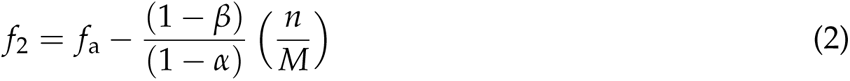

Since *α < β*, it is clear that *f*_1_ *< f*_2_; the beneficial fraction for mutations in the over-sampled class is less than the beneficial fraction in the under-sampled class.

We can also compare the two DFEs defined above: (1) DFE_all_, the DFE of all possible mutations that assumes equal access to all mutations, and (2) DFE*_β_*, the mutation-weighted DFE that takes the bias into account. Let *f*_all_ and *f_β_* denote the fractions of beneficial mutations in DFE_all_ and DFE*_β_*, respectively, after this period of evolution. Since DFE_all_ samples class 1 mutations with probability *α* and class 2 with probability (1 *− α*), we find

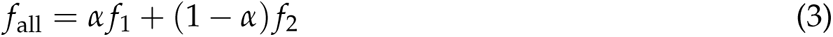

while similarly

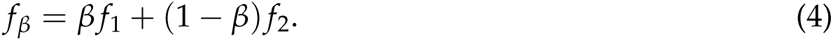

Again, since *α < β* and *f*_1_ < *f*_2_, we find that *f*_all_ > *f_β_*. Hence, the organism will access a smaller fraction of beneficial mutations than is potentially available.

This might change, however, if the mutation bias shifts. If the mutation bias shifts to *β^′^*, then the bias-weighted DFE changes and the fraction of beneficial mutations becomes 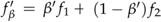. There are two possibilities: (1) the new bias reduces or reverses the previous bias. Taking as we did before the case when *α < β*, a bias reduction occurs when *α < β^′^ < β*, while a reversal happens when *β^′^* is even lower that *α*. In either case (reduction or reversal), we find that 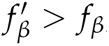, in other words, the new bias increases access to beneficial mutations. In contrast, (2) the new bias could reinforce the original bias, which occurs when *β^′^ > β*. This yields 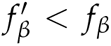 and thus implies reduced access to beneficial mutations. Moreover, it is clear that these effects will be stronger when the difference *|β − β^′^|* is bigger, i.e., when the reversal or the reinforcement has a greater magnitude.

In the simulations to follow, we illustrate examples of these effects where mutations are classified as transitions and transversions, and in which we relax the assumptions of a smooth fitness landscape (no epistasis), of rare beneficial mutations, and of strong-selection weak-mutation evolutionary dynamics.

### Hitchhiking probabilities

Some simple “back of the envelope” calculations offer critical insights into the evolution of the mutation bias. We follow a heuristic argument described by Lenski (2004) and developed more formally by Wylie et al. (2009). Suppose a fraction *p_µ_* of a population has an *F*-fold higher mutation rate, but is otherwise identical to the wildtype. The chance that the next beneficial mutation that sweeps through the population occurs in the mutator strain is then simply approximated by *p_µ_F*. Lenski gives a quantitative estimate of this factor for *E. coli* mutators that segregate at an estimated frequency of 1 *×* 10^−4^ and have a 100-fold increase in mutation rate, predicting that about 1% of beneficial substitutions will occur in the mutator.

This argument naturally assumes that the DFE is identical in the wildtype and mutator. In contrast, suppose a fraction *p*_bs_ of a population has a shift in mutation bias, but has the same mutation rate as the wildtype. If the bias shift reduces or reverses the wildtype bias, the bias-shifted strain will experience a *G*-fold higher beneficial fraction:

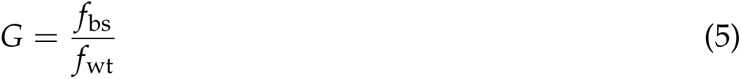

where *f*_wt_ and *f* bs denote the beneficial fraction of the DFE*_β_* in the wildtype and bias-shifted strain respectively. Assuming the effect size of beneficial mutations and the deleterious load are unchanged with the bias shift, the chance that the next beneficial mutation carries the bias shifted strain to fixation is approximated by *p*_bs_*G*.

If, in addition, the beneficial effect size is increased in the bias-shifted strain, then the expected fixation probability of a beneficial mutation in this strain, *π*_bs_, will exceed the analogous expectation in the wildtype, *π*_wt_, such that beneficial mutations have an *H*-fold higher fixation probability if they occur in the bias-shifted background, where:

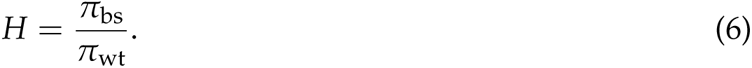

Overall, the fraction of beneficial fixations that occur in the bias-shifted strain is then *p*_bs_*GH*.

A quantitative estimate of this factor is challenging. In bacteria, both environmental stress and the loss of function of specific DNA repair enzymes can cause bias shifts without a substantial change in mutation rate (Foster et al., 2015; Maharjan and Ferenci, 2017; Sane et al., 2021; Shewaramani et al., 2017), however it is difficult to estimate the frequency at which such mutations might segregate in natural populations. Nonetheless, in both experimental (Sane et al., 2021) and simulated fitness landscapes as seen in the Results to follow, both factors *G* and *H* (the relative advantage of bias-shifted strains) are modest, typically in the range of one to two. In contrast, *F*, the mutation rate multiplier, can be a factor of 100 or more. We conclude that unless bias-shifted strains are maintained by the mutation-selection balance at frequencies that are several orders of magnitude higher than mutator strains, they will be much less likely than mutators to fix during adaptation.

However, suppose a fraction *p_µ_*_,bs_ of a population has *both* an increased mutation rate and a bias-shift. This is in fact the most likely situation for loss-of-function mutations in DNA repair (Foster et al., 2015; Sane et al., 2021). The chance that a beneficial mutation carries this strain to fixation would then be *p_µ_*_,bs_*FGH*, in other words, the effects combine multiplicatively. We thus predict, in agreement with Couce et al. (2013), that bias shifts could powerfully affect the invasion rate of mutator strains.

The reasoning above neglects the effects of changes to the deleterious load when the bias or mutation rate is altered. Recall that for a mutation rate *µ* per genome and deleterious fraction *d* in DFE*_β_*, the load is approximated by the product *µd*, and is in fact independent of the deleterious effect size (Ewens, 2012). Thus a strain that increases the mutation rate only, by factor *F*, will experience an increased load of *Fµd*. In contrast, for strains in which only a bias shift occurs, the deleterious fraction, *d*, will be reduced whenever the beneficial fraction *f* is increased (although this reduction is exactly concomitant only in very large populations in which the neutral fraction of the DFE is negligible). Thus any bias shift that increases the beneficial fraction will reduce the load.

For a mutator strain in which both the mutation rate and bias are altered, both of these effects come into play. Depending on the relative magnitudes of the increase in mutation rate and reduction in the deleterious fraction, the mutator strain may carry a greater or lesser load than the wildtype.

The magnitude of deleterious load will ultimately affect the fixation probability for beneficial mutations on the mutator background, reducing (and possibly reversing) the fixation advantage. For a mutator with an increased mutation rate *Fµ*, load overwhelms the expected selective advantage of new beneficial mutations when *Fµd > s*_b_, or when the mutation rate multiple exceeds the critical value

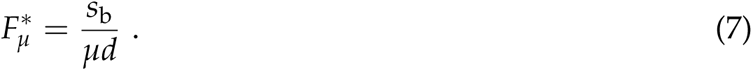

In constrast, consider a bias shift that increases the expected beneficial effect by a factor (1 + *δ*), and also reduces the deleterious fraction by a factor (1 *− ɛ*). For a mutator in which both the bias and mutation rate change, load overwhelms the selective advantage when *Fµd*(1 *− ɛ*) > *s*_b_(1 + δ), or when *F* exceeds

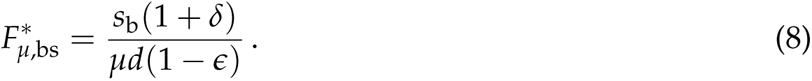

Since 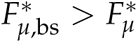, these results predict that bias reductions and reversals will not only increase the invasion probability of mutator strains, but will extend the range of mutation rates over which invasion is favoured. Both of these predictions are verified in the simulation results to follow.

### Simulation Methods

We simulated population evolution on the well-studied *NK* fitness landscape (Macken and Perelson, 1989; Stoltzfus, 2006). Each genotype is represented by a genomic sequence of length *N*, composed of four bases (A, C, G, T) such that mutations can be classified as transitions or transversions.

In an *NK* fitness landscape, the fitness of a sequence is simply calculated by adding the fitness contributions of all its loci. To incorporate epistasis, however, the fitness contribution of each locus depends not only on the state of the locus, but on the states of *K* other randomly assigned loci. Thus *K* determines the degree of epistasis, i.e., the number of loci epistatically coupled to each locus. For instance, for *K* = 2 and a given locus *m*, there are 4*^K^*^+1^ = 64 different possible combinations for that locus and its two neighbours: AAA, AAC, AAG, TTT. Every such combination is randomly assigned a uniformly distributed value from the interval (0, 1) which defines the fitness contribution of locus *m*.

Each individual in the population is assigned the following: (1) a genome sequence, which determines fitness; (2) a transition:transversion bias *Ti* : *Tv*, where *Ti* = 1 *− Tv* gives the probability that a *de novo* mutation is a transition; and (3) a mutation rate *µ* per genome per generation. In some simulations the bias and/or mutation rate are fixed, while in other scenarios they are manipulated.

The population is initialized in generation 0 with a population size *n*_0_, typically seeded with a number of random genotypes. Generations are discrete; the number of offspring for the *i*th genotype in generation *j* is Poisson distributed with mean

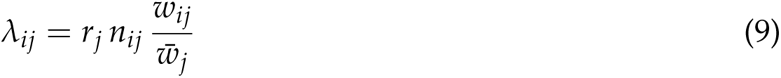

where *n_ij_*and *w_ij_*represent the number of individuals and the fitness of the *i*th genotype in generation *j*, respectively, while 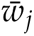 is the mean fitness of the population in generation *j*. The variable 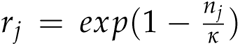 is the Ricker factor in generation *j*, which limits overall population growth by comparing the total number of individuals in generation *j*, *n_j_*, to the carrying capacity *κ*.

In every generation, mutations occur according to each genotype’s mutation rate, *µ_i_*. When a mutation occurs, it affects a randomly chosen nucleotide in the sequence, and the mutation is a transition or a transversion according to that genotype’s bias. If the mutation is a transversion, one of the two possibilities (e.g. *T → G* or *T → A*) is chosen at random. Mutations occur during reproduction and when they occur, a single individual with the new genotype is added to the next generation, where the new individual inherits the parental mutation rate and bias.

We keep track of each distinct genotype in the population, and number of individuals of that genotype, but we do not track the ancestry of every individual. Thus in order to estimate – at a given generation – the divergence of the population from the initial population, we first identify the most common genotype in the population. We then compute the Hamming distance (number of nucleotide differences) between this genotype and each genotype in the founding population. We report the minimum of these Hamming distances as a measure of genetic divergence from the ancestral population. As a proxy for “fixations”, we also report the number of transitions and transversions by which the most common genotype differs from its closest match in the founding population. This is a proxy for two reasons: because true “fixations” may or may not occur in our genetically diverse full population simulations; and also because two consecutive transversions at the same locus might lead to what would be counted as a transition.

In the results to follow, we perform invasion tests for mutant strains that change only the mutation bias but not the mutation rate, mutators that increase the mutation rate alone, and mutators that change both the rate and bias, as illustrated in figure 1. Invasion tests are initiated in our simulations either by following the fate of in an invading subpopulation at an initial frequency of 5%, or by tracking the fate of a single randomly chosen individual. Because any single lineage has a high probability of going extinct even if it carries a selective advantage, the latter approach may require simulating a very large number of replicates. Thus, for computational efficiency, results in the main text were generated for an invading subpopulation. (Figure S1 shows results for the invasion of a single individual; we observed no qualitative differences in invasion results in the two cases.) We also note that predicted outcomes with an invader frequency of 5% are amenable to empirical testing.

**Figure 1:**
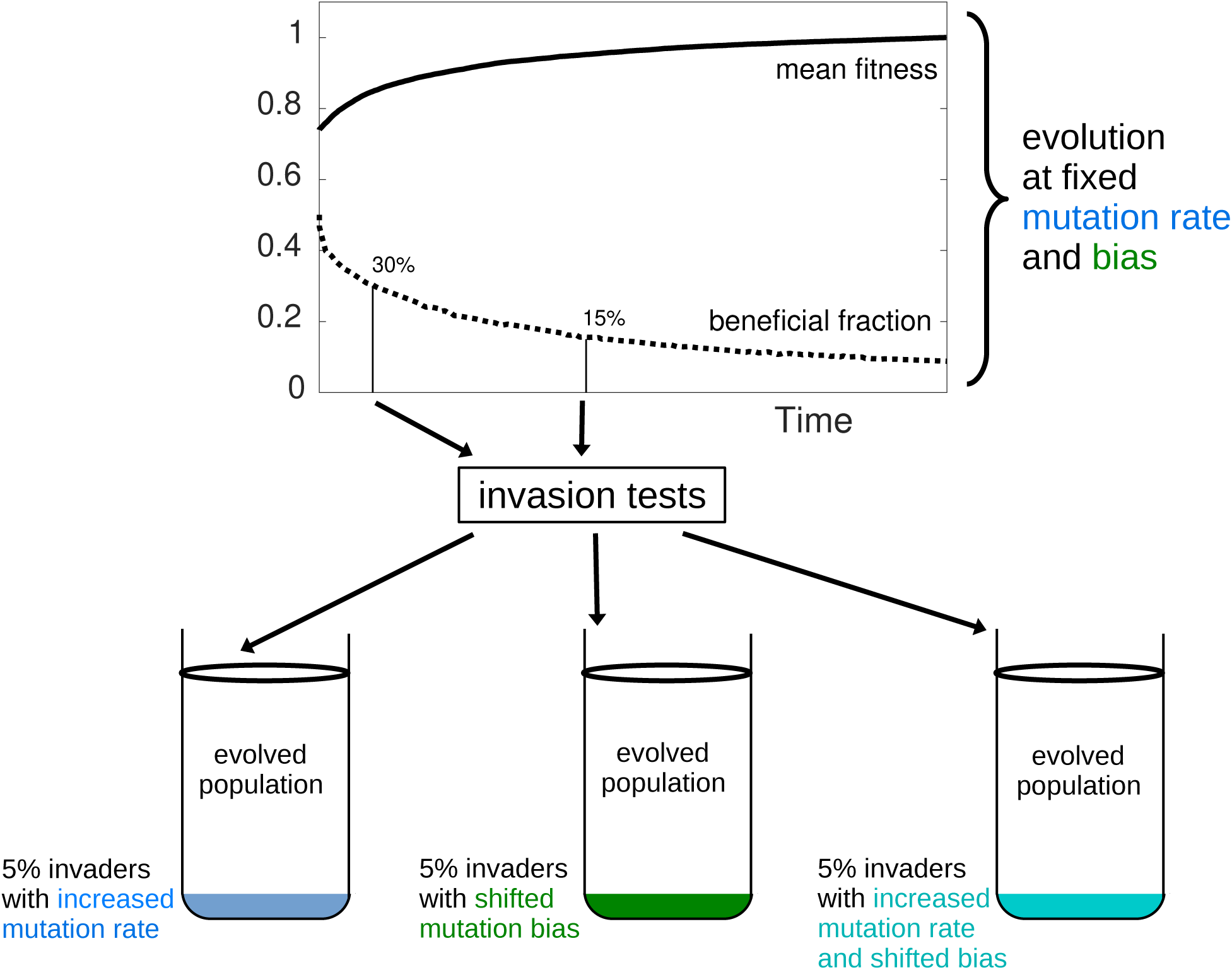
Schematic diagram of simulated invasion tests. Populations of initially random genotypes evolve with a fixed mutation rate and fixed transition:transversion bias. The mean population fitness (solid line) increases over time, while the fraction of mutations in the mutation-weighted DFE that are beneficial (dotted line) declines. When the mutation-weighted beneficial fraction, *f_β_*, reaches 30%, we perform the three invasion tests illustrated. The same tests are repeated in new simulations but when the beneficial fraction in the evolving population reaches 15%. For each ‘replicate’, we create a new fitness landscape, initiate a new population, and evolve starting at time zero.

Populations in these simulations, however, are genetically diverse; in some parameter regimes the average number of segregating strains is of order 100. The use of an invading subpopulation thus necessitates the choice of a genotype for the invading strain. Since the most common genotype in the population is typically also the fittest, using this genotype for the invading strain biases the results in favour of invasion. Instead, we initiate the mutant subpopulation by changing the mutation rate and/or bias in a random 5% of individuals in the population. Thus, the invading sub-population has on average the same genetic diversity (and, critically, the same genetic load) as the ancestral population.

Similar to a competition experiment, we then track whether this sub-population invades. We perform invasion tests in populations at various degrees of adaptive potential, which we estimate based on the remaining fraction of beneficial mutations in the DFE*_β_* for the most common genotype. The invasion probability is estimated as the fraction of *n*_reps_ replicates in which a lineage from the original subpopulation becomes the most common genotype in the population. For an estimated invasion probability *p*, the precision of the estimate is given by the standard deviation of a binomial random variable: 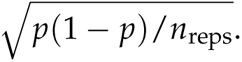

### Parameter values

Unless otherwise indicated, our simulations used a sequence of length *N* = 100 and epistasis parameter *K* = 0 (no epistasis) or *K* = 1 or 2. We used a carrying capacity of *κ* = 5000 and initialized simulations with 50 random genotypes, each with *n_i_*_,0_ = 100 individuals. We note that for a randomly generated genotype in the NK-model, 50% of mutations are expected to be beneficial.

The default sequence mutation rate in the simulations was *µ* = 0.0001 per genome per generation, the order of magnitude of *E. coli* K12 (Sane et al., 2021). Invasion success is determined after 5000 generations. Note that our results were not sensitive to this final time, as long as it was sufficiently large. See Table 3 for a list of simulation parameters and default values.

**Table 3:** Simulation parameters and their default values.

| Parameter | Symbol | Default Value |
| --- | --- | --- |
| Genome length | $N$ | 100 |
| Epistasis degree | $K$ | 0, 1, or 2 |
| Mutation rate (per genome per generation) | $\mu$ | $10^{-4}$ |
| Initial number of genotypes |  | 50 |
| Initial population size for genotype $i$ | $n_{i0}$ | 100 |
| Carrying capacity | $\kappa$ | 5000 |

## Results

### Adaptive substitutions reflect the mutational bias

Using differences between the most common genotype and its closest genetic relative in the founding population as a proxy for fixation, figure 2(a) demonstrates that the mutation bias is strongly reflected in the substitutions that occur as the population adapts. In the illustrated case, a bias of *β* = 0.7 (70% transition probability) results in the number of fixed transitions exceeding fixed transversions, even though there are twice as many available transversions as transitions.

**Figure 2:**
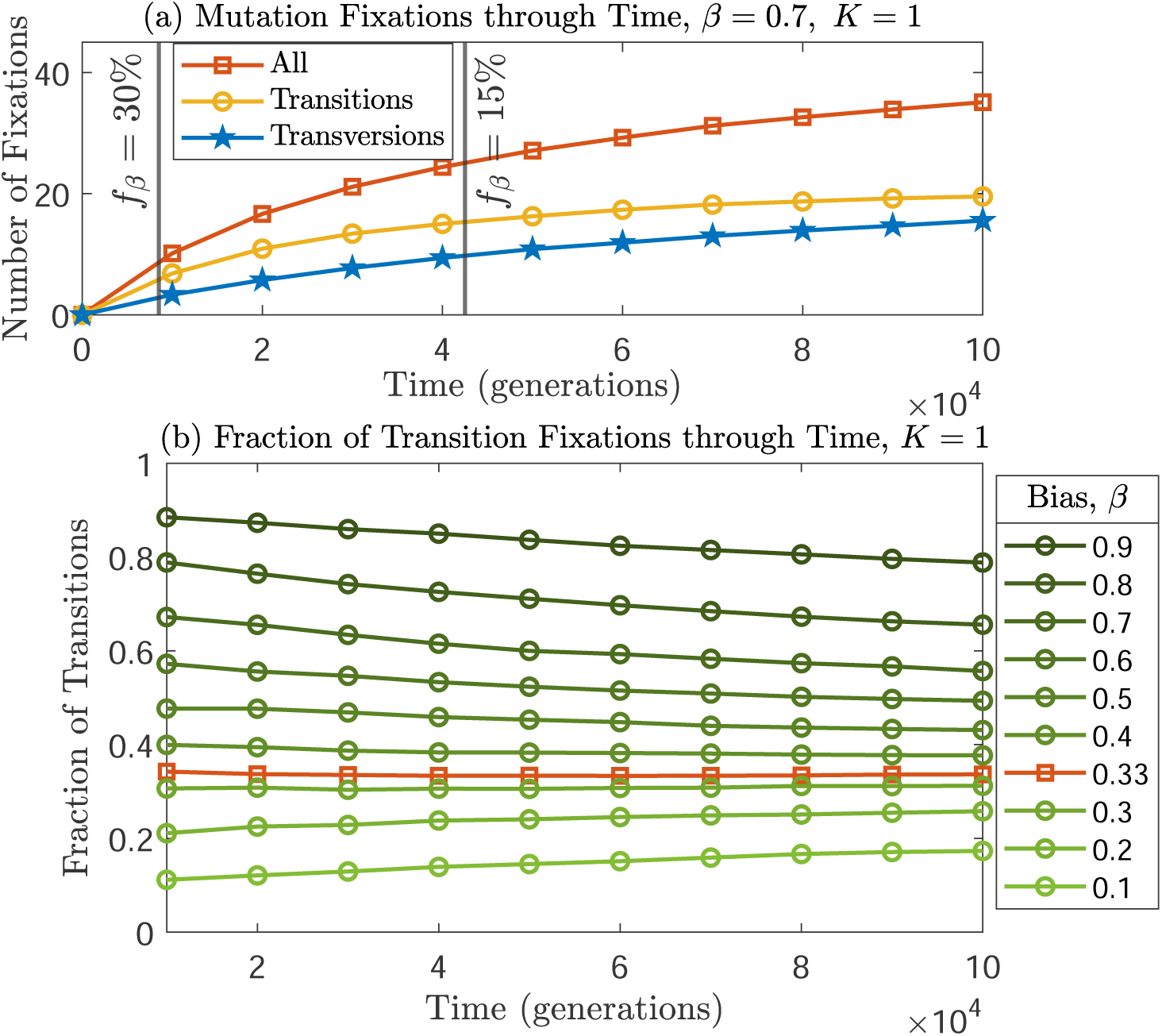
The numbers of fixations of transitions and transversions through time are affected by the mutational bias. (a) More transitions (yellow circles) than transversions (blue stars) fix in a transition-biased population with *β* = 0.7 and an epistasis degree of *K* = 1. The Hamming distance to the closest ancestor, the sum of transition and transversion fixations, is shown in red squares. Two vertical lines are added to clarify the times at which the bias-weighted beneficial fraction, *f_β_*, reaches 30% and 15% (b) The fraction of fixations that are transitions is almost constant for unbiased mutations with *β* = 1/3 (red squares), while it strongly reflects the mutation bias in other cases. As adaptation proceeds, this fraction slightly decreases for transition-biased populations (above the red squares) and slightly increases for transversion-biased populations (below the red squares). The standard error of means in the relative fixations is high for the first few thousand generations due to the low number of fixations (on average across all biases, less than 1 mutation fixes at generation 1000, while only 5.5 mutations fix at 5000 generations). Thus, results are shown starting at generation 10,000 for panel (b). Means of 300 replicates are shown. Error bars are smaller than symbol heights and are omitted.

In figure 2(b), we plot the fraction of fixations that are transitions, versus time, for different values of the mutation bias. It is clear from the figure that substitutions reflect the bias strongly, particularly at early times. As adaptation proceeds, the over-represented class fixes relatively less often as beneficial mutations of that class are depleted. If mutation is unbiased, *β* = 1/3, one-third of substitutions are transitions as expected, and this value remains constant.

### The fraction of beneficial mutations in the over-sampled mutation class rapidly declines

As beneficial mutations in over-sampled mutation classes fix more frequently than those in under-sampled classes, our analytical work predicts that this reduces the beneficial fraction of the DFE for over-sampled classes. We first confirm this effect in full populations by simulating the evolution of either transition- or transversion-biased populations, recording the beneficial fraction of all possible transitions, *f*_Ti_, and all possible transversions, *f*_Tv_, for the most common genotype in the population at any time.

Figure 3 shows results for a a transition-biased population with *β* = 0.7, consistent with a two-fold excess of transitions above the null expectation (Stoltzfus and McCandlish, 2017). Although the beneficial fraction is initially the same in both mutational classes, *f*_Ti_ decays more quickly than *f*_Tv_ (fig. 3(a)). The overall beneficial fraction declines but remains between *f*_Ti_ and *f*_Tv_. The difference between *f*_Ti_ and *f*_Tv_ increases with time as beneficial transitions fix more often than beneficial transversions, and is significantly different from zero except at very early time points (panel (b)). After 100,000 generations, there are roughly twice as many beneficial transversions as transitions available (panel (b)); this factor of two also reflects the maximum possible increase in the beneficial fraction for a bias-shifted strain, equivalent to the factor *G* in the heuristic argument above, for an extreme bias shift from transitions to transversions. Although this factor could increase further at later times, the simulated populations have little remaining adaptive potential at the end of the simulation time.

**Figure 3:**
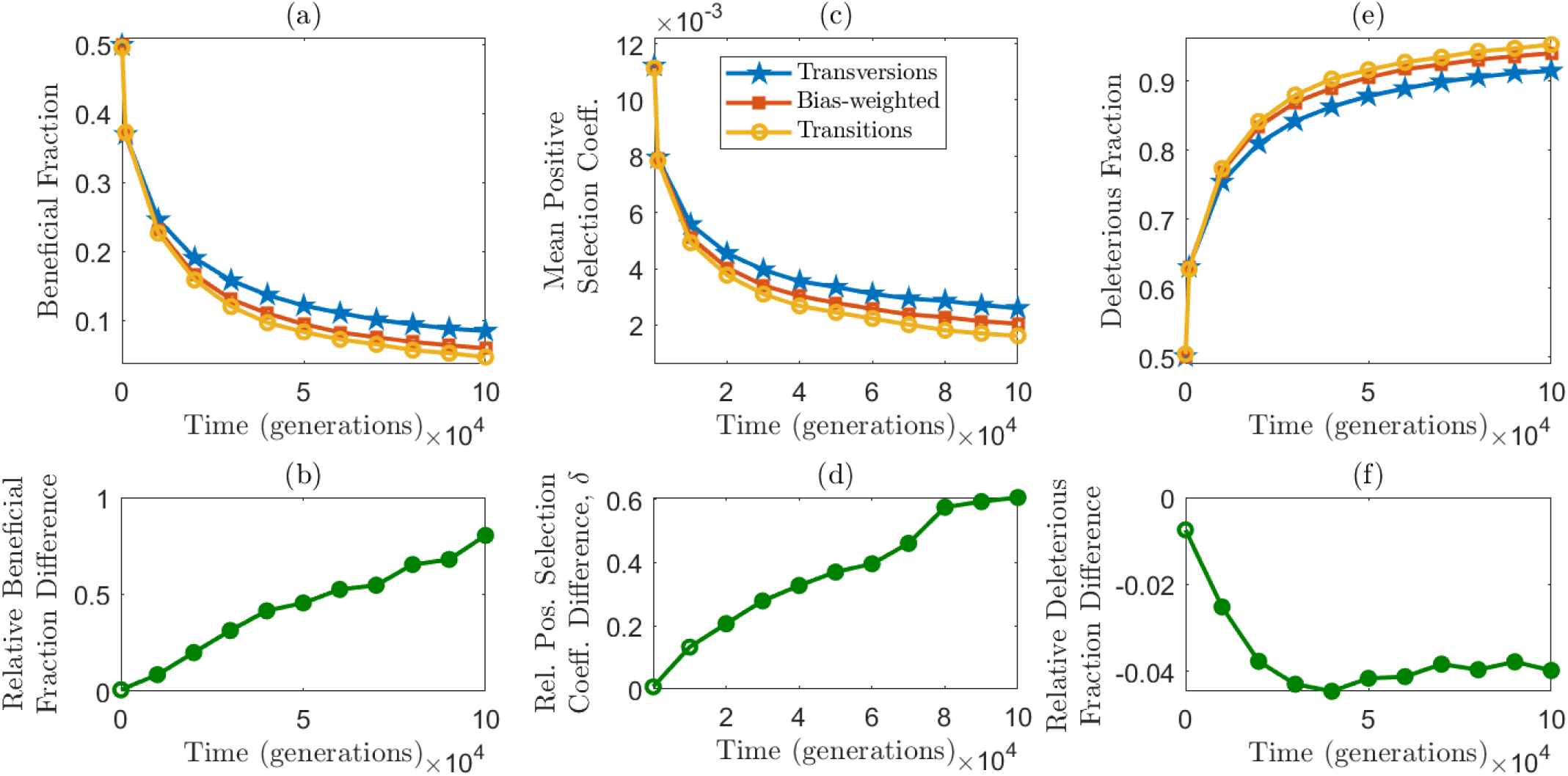
Beneficial fractions and mean beneficial effect sizes decline during the evolution of a transition-biased population, while deleterious fractions increase. (a) The fraction of beneficial transitions (the over-sampled class, yellow circles) decreases over time, falling more rapidly than the fraction of beneficial transversions (blue stars). The overall beneficial fraction of the bias-weighted DFE ( *f_β_*, red squares) falls between these two extremes (closer to *f*_Ti_ as expected since the transition frequency *β* = 0.7). (b) As a result of their different decay rates, the relative difference in the beneficial fraction, ( *f*_Tv_ *− f*_Ti_)/ *f*_Ti_ increases over time; filled circles indicate that the difference *f*_Tv_ *− f*_Ti_ is significantly different from zero at that time (t test, *p <* .05). Panels (c) and (d) show analogous results for the mean positive selection coefficients of transitions and transversions. (e) As the beneficial fraction decreases, the deleterious fraction increases. The fraction of deleterious transitions (yellow circles) increases more rapidly than the fraction of deleterious transversions (blue stars). (f) The relative difference in the deleterious fraction increases in magnitude over time, but the effect is modest (less than 5% difference). In all cases, means of 300 simulation replicates are shown for *β* = 0.7 and epistasis degree *K* = 2; DFEs are computed for the most common genotype in the population at each time point. Error bars in top panels are smaller than symbol heights and are omitted.

In addition, transversions not only exceed transitions in their beneficial fraction, but also in the mean magnitude of the positive selection coefficient; in other words, as evolution progresses a single beneficial transversion is expected to be more advantageous than a single beneficial transition (fig. 3(c) and (d)). At the end of the simulation time, the effect size of a beneficial transversion was about 1.5-fold higher than a beneficial transition, corresponding to factor *H* in the heuristic argument.

As evolution proceeds and the beneficial fraction of the DFE is reduced, the deleterious fraction concomitantly increases. Again, the deleterious fraction for both mutational classes is initially 50%, but the deleterious fraction for the oversampled class, *d*_Ti_, increases more quickly than *d*_Tv_ (fig. 3(e)). The difference between *d*_Ti_ and *d*_Tv_ increases in magnitude over time (panel f), but the effect is more modest than the changes observed in the beneficial fraction. After 100,000 generations, the difference in the deleterious fraction is less than 5%. Overall, the observed increase in the deleterious fraction predicts a modest increase in the deleterious load for an over-sampled mutational class. We note for completeness that the mean negative selection coefficient changes only negligibly during evolution (see fig. S2); differences between the deleterious selective effect for transitions and transversions were at most 2%.

The qualitative behaviours observed in figure 3 were robust across the parameter ranges we tested, including smooth fitness landscapes (fig. S3, *K* = 0) and for a lower degree of epistasis (fig. S4, *K* = 1). The magnitude of these effects is reduced for weaker transition biases, (fig. S5, *β* = 0.5), and is reversed when the population is initially transversion-biased (fig. S6, *β* = 0.1).

### Bias reversals increase the invasion probability of mutators, and can reverse the expected outcome of invasion

Given the potential increase in beneficial fraction afforded by a bias shift, we next asked whether strains with a bias shift would invade a wildtype population. Figure 4 shows results for both smooth (blue) and epistatic (red) fitness landscapes, for populations at two degrees of adaptive potential (30% and 15% remaining beneficial mutations, left and right panels respectively), and for the limiting case of a full bias shift (*β* = 1 *→ β^′^* = 0, top panels) and a second case of a strong bias shift (*β* = 0.9 *→ β^′^* = 0.1, lower panels).

**Figure 4:**
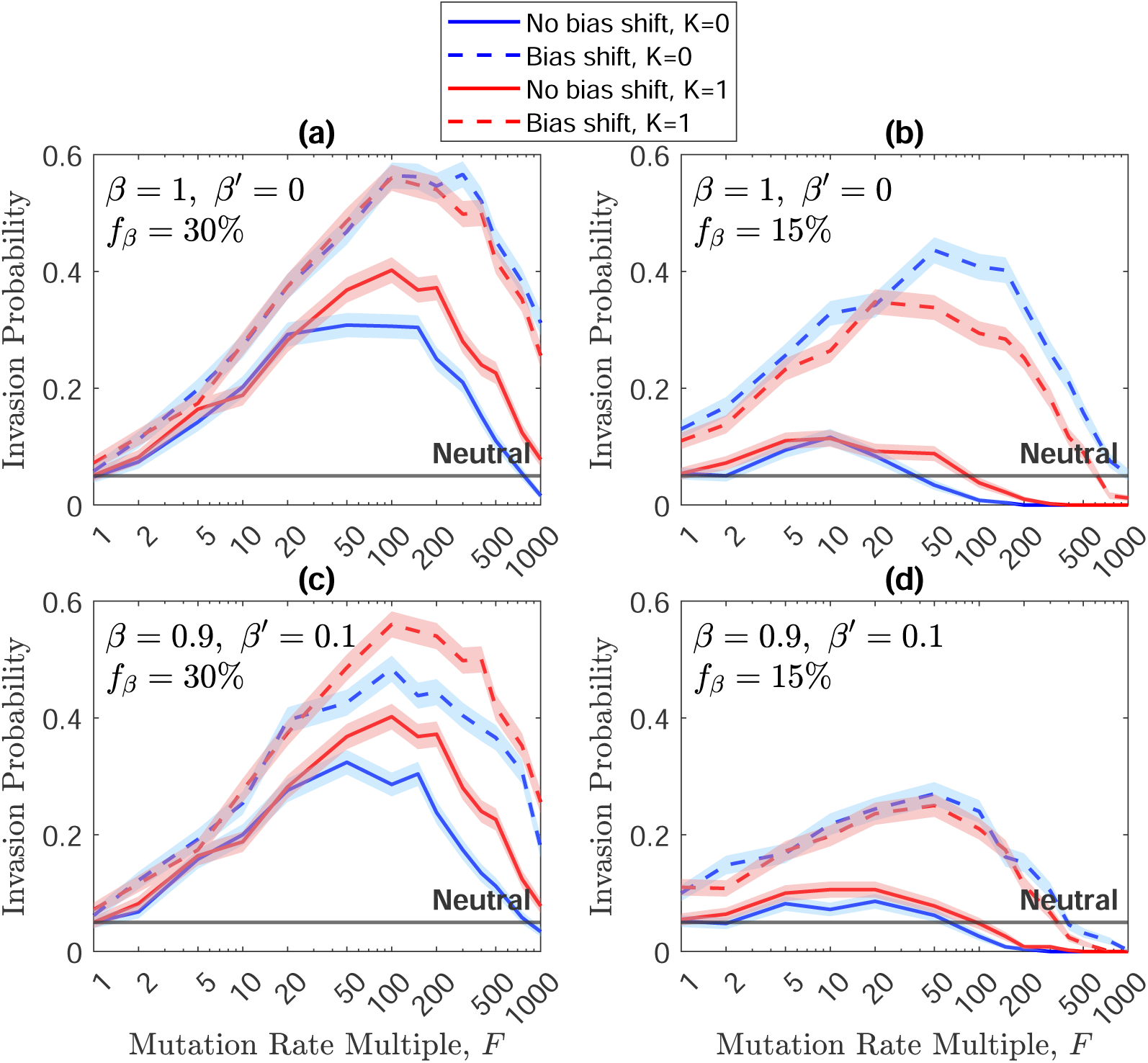
Invasion probabilities for strains with both bias shifts and increased mutation rates (dashed lines) exceed invasion probabilities for strains that increase the mutation rate only (solid lines). Simulation results are shown in which each invading strain is initiated at 5% of the population and has an *F*-fold increase in the mutation rate (*x*-axis); we compare no change in bias (solid lines), with a full change in bias (*β* = 1 *→ β^′^* = 0, top row), or a strong change in bias (*β* = 0.9 *→ β^′^* = 0.1, bottom row). Invasion tests occur for populations at two levels of adaptive potential, *f_β_* = 30% (left panels) and *f_β_* = 15% (right panels). The case of epistasis (*K* = 1) is shown in red, while the blue lines show the case of no epistasis (*K* = 0). The horizontal lines represent the neutral expectation 0.05, which represents the invasion probability of a 5% subpopulation in the absence of phenotypic change. Results of 500 replicates are shown; shaded regions indicate *±* one standard deviation.

As a comparison, we tested the invasion probability of mutator strains with an *F*-fold increase in mutation rate (*x*-axis), but no change in the mutation spectrum (solid lines). On the left edge of each panel (*y*-intercepts), we confirm that strains that change neither the mutation rate nor the bias have a 5% invasion probability when seeded at an initial frequency of 5%, the neutral expectation. As the mutation rate multiple, *F*, is increased, mutator strains are initially favoured (have an invasion probability higher than neutral), particularly in populations with more adaptive potential (left panels). Further increases in mutation rate become disfavoured, however, as the deleterious load overwhelms the benefit of accessing beneficial mutations more easily. In general we see that mutators are strongly favoured when many beneficial mutations are available (left panels), but high mutation rates are disadvantageous when the population is closer to the fitness peak (right panels). We also note that mutators are favoured on epistatic landscapes (solid red versus solid blue), presumably because of an enhanced ability to cross fitness valleys on rugged landscapes.

We then tested shifts in mutation bias (dashed lines). The left edge (y-intercept) of each panel thus shows the invasion probability for a strain that has a bias shift (dashed lines) but no change in mutation rate; in all cases the bias shift increases the invasion probability above the neutral expectation, but even in the most extreme case ( *f_β_* = 15% and *β* = 1 *→ β^′^* = 0), the advantage of the bias-shifted strain is rather modest.

The effect of a bias shift can be dramatically increased, however, for mutator strains (dashed lines versus solid lines at *F >* 1). In all cases, mutators have substantially higher invasion probabilities if they also reverse the mutation bias. As predicted theoretically, in all cases the bias shift extends the range of mutation rates over which mutators are favoured. For mutator strains with very high mutation rates, the invasion probability can change from below the neutral expectation or near zero, without the bias shift, to values as high as 20-40%. We also point out that the effect of combining an increase in mutation rate with a bias shift, in these full population simulations, is highly nonlinear. If we take as an example the smooth fitness landscape (blue lines) in figure 4(b), a 100-fold increase in mutation rate, alone, reduces the invasion probability below the neutral expectation, while the bias shift, alone, increases the invasion probability by a factor of just over two (*y*-intercept). However a strain that both shifts the bias and increases the mutation rate 100-fold has an invasion probability that is 8-fold higher than neutral.

### The direction of the bias shift relative to an unbiased spectrum determines the effect of the shift

Having established that bias shifts, at least in these extreme cases (1 *→* 0 and 0.9 *→* 0.1), are more likely to emerge when coupled with changes in mutation rate, we now investigate bias shifts at a range of directions and magnitudes on the fate of mutator strains.

As described in the *Theory* section, the effect of a bias shift depends on the unbiased mutation frequency, *α*, the bias with which the population has been evolving prior to the shift, *β*, and the shifted bias *β^′^*. Figure 5 shows how this applies to the transition-transversion ratio, where *α* = 1/3. We first investigate changes in the pre-existing bias, *β*. In particular, after populations have evolved with various values of *β*, we quantify the invasion probability of a mutator strain with a 50-fold increase in mutation rate, comparing the fate of this mutator strain without a bias shift (solid lines) to its fate with a bias shift (dashed lines). Results for two values of *β^′^* are shown in the two panels in figure 5. Panel (a) shows that, when the population is initially transition-biased (to the right of the vertical line), a shift to *β^′^* = 0.9 reinforces the previous bias, and the invasion probability is reduced by the bias shift; in contrast, if the population is initially transversion-biased (to the left of the vertical line), a bias shift to *β^′^* = 0.9 reverses the existing bias, and increases the invasion probability. These two effects are reversed when the bias shifts to *β^′^* = 0.1, as shown in panel (b). (Results for reinforcements in (b) are not significantly different with and without bias shifts, presumably due to the limited possible magnitude of bias shifts in this region.) Results are shown for both smooth and epistatic fitness landscapes, and for populations at an adaptive potential of 15% remaining beneficial mutations. Similar results are obtained when the mutators have a 10-fold higher mutation rate (fig. S7), and also at an earlier time when the beneficial fraction is 30%, either for the same mutation rate multiple of *F* = 50 (fig. S8) or for *F* = 200 (fig. S9). The advantage of a bias reversal is more striking when *f_β_* = 15%; at this stage the DFE for the ancestor is depleted and the bias shift becomes a key factor in the competition between ancestor and mutator.

**Figure 5:**
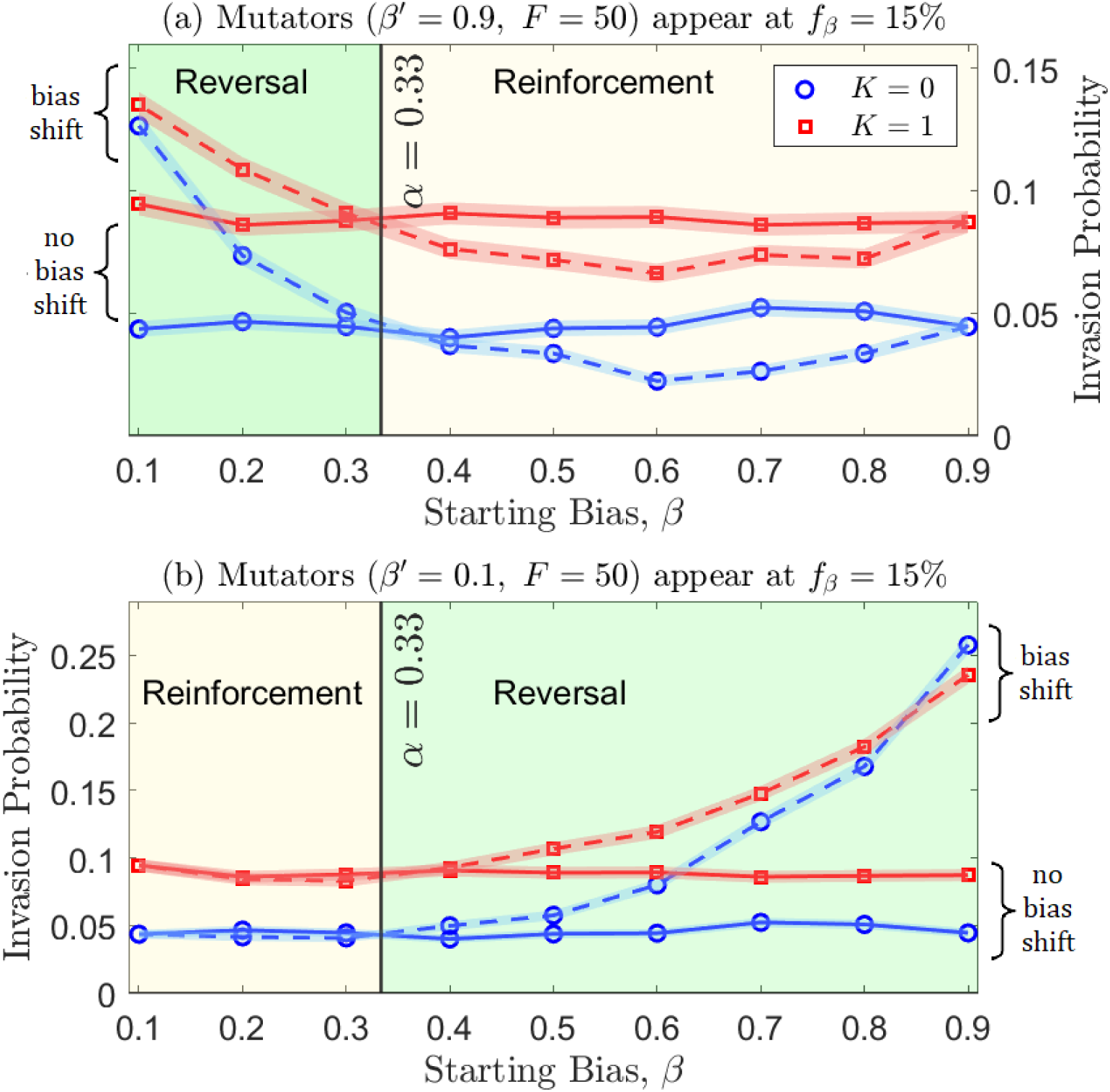
Invasion probabilities of mutators increase when the bias is reversed. Populations evolved with a fixed starting bias (*x*-axis) until the beneficial fraction of the most common genotype reached 15%. An invasion test was then simulated for a strain with a 50-fold increase in mutation rate. Two cases are compared: the mutators either have the same bias as the initial population (solid lines), or have a bias shift to (a) *β^′^* = 0.9 or (b) *β^′^* = 0.1 (dashed lines). The vertical line at *α* = 1/3 represents the unbiased transition frequency. Panel (a) shows that for a starting bias less than *α*, a shift to *β^′^* = 0.9 reverses the bias and thus increases the invasion probability (dashed lines above solid lines); the same bias shift tends to reduce the invasion probability (dashed lines below solid lines) if the starting bias is greater than *α*. The effects are reversed when the bias shifts to *β^′^* = 0.1, as shown in panel (b). Results of 4000 replicates are shown; shaded regions indicate *±* one standard deviation.

### Stronger bias reversals increase the invasion probability

While the direction of the bias shift establishes whether it is beneficial, the magnitude of the shift determines the magnitude of the benefit. Figure 6 shows that stronger reversals have a higher invasion probability, again in the presence or absence of epistasis and for populations relatively early and later in adaptation. In contrast with figure 5, here we fix the starting bias, *β* = 0.9, and shift the bias to various values of *β^′^*. Thus bias reversals occur when the new bias *β^′^* is smaller than the unbiased value *α* = 1/3; in this case the spectrum changes from transition- to transversion-biased. Bias reductions occur when *β^′^ < β* but *β^′^ > α*; the new spectrum is still transition-biased, but is less extreme. When *β^′^ >* 0.9, the bias is reinforced. We note that the influence of the bias shift is greater (steeper slope in figure 6(a)) at *f_β_* = 15% than at *f_β_* = 30%, again presumably because the bias shift is less critical to ”finding” a beneficial mutation when there are more beneficial mutations available. Invasions are tested for mutator strains with a 50-fold increase in the mutation rate in figure 6(a), whereas in figure 6(b) tests are performed for mutation rate multiples *F* = 10, 50, 200. All of these simulations give the same qualitative results, illustrating the advantage of reducing or reversing a prevailing bias.

**Figure 6:**
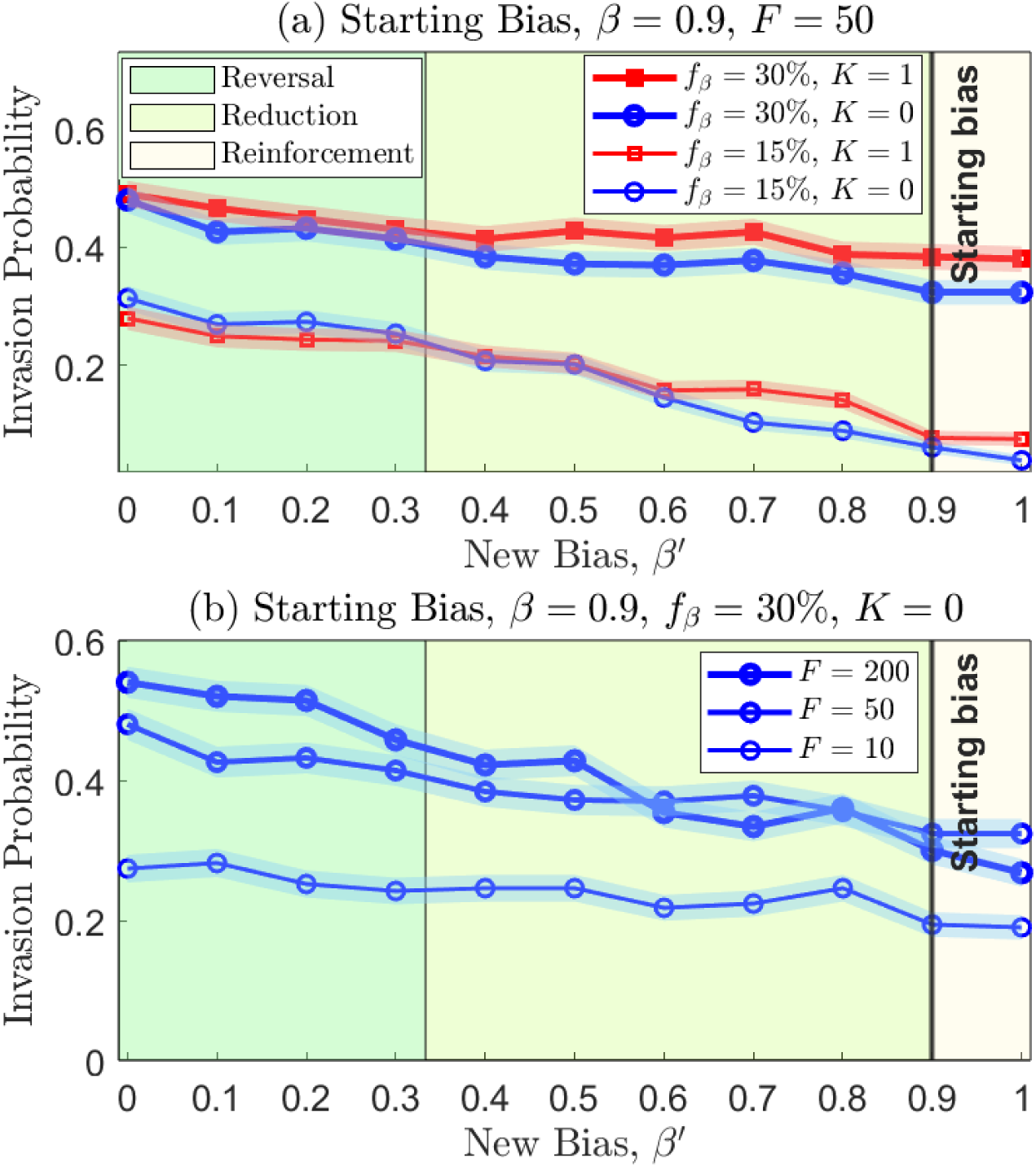
In populations that have evolved with a transition frequency of *β* = 0.9, the invasion probability of mutators is increased when the bias is reduced or reversed. (a) Invasion tests were initialized for a mutator strain with a 50-fold increase in mutation rate and a bias shift to *β^′^* as indicated (*x*-axis). Larger reversals increased the invasion probability whether the invasion test occurred at a bias-weighted beneficial fraction of 30% (bold lines) or 15% (thin lines), on either smooth (*K* = 0, blue circles) or epistasic landscapes (*K* = 1, red squares). (b) Results are consistent across different values of the mutation rate multiplier, *F* (*K* = 0, *f_β_* = 30% illustrated.) Results of 500 replicates are shown; shaded regions indicate *±* one standard deviation.

## Discussion

Multiple factors can affect the evolution of the mutation rate and spectrum. Population genetics theory predicts that for a given number of functional sites in the genome, the larger the effective population size the greater the power of natural selection to reduce the mutation rate (drift-barrier hypothesis, (Lynch, 2011; Sung et al., 2012)). In animals, selection on sequence-dependent DNA repair enzymes, which could potentially alter the mutation spectrum within a genome, is likely to fall below the threshold at which natural selection is effective (Harris and Pritchard, 2017). Hence, there is the possibility that in natural populations DNA sequence-dependent hypo/hyper-mutators could segregate. For example, a natural hypomutator allele reducing the mutation rate of C*→*A mutations has been recently discovered in mice (Sasani et al., 2022). In humans, the TCC*→*TTC mutation rate increased in Europeans 15,000 to 2,000 years ago (Harris and Pritchard, 2017), but it is unknown whether this was driven by an environmental mutagen or a hypermutator allele that temporarily rose in frequency. In fungi, it has been recently described how the presence/absence of 52 DNA mismatch repair (MMR) genes impacts the mutation spectrum across more than 1,000 species, finding that in pathogenic species the loss of MMR genes is substantial and has probably increased their mutation rate (Phillips et al., 2021). In human tumors, mutations on MMR genes have a characteristic signature in the mutation spectrum (Alexandrov et al., 2013) that can be recovered in *C. elegans* when mutating the same genes (Meier et al., 2018). Other DNA repair pathways and genomic integrity checkpoints, such as Rad53p (human homolog Chk2), have been described to be down-regulated in fungal pathogens (Shor et al., 2020; Steenwyk, 2021). These empirical works highlight the complex interaction between population size and the environment as drivers of mutation rate and spectrum evolution, and possibly also mutation bias evolution. Finally, apart from the number of functional sites in the genome, other genomic attributes such as the recombination rate can also impact the efficacy of natural selection on mutation rate modifiers (sequence-context dependent or not). Increasing the recombination rate is expected to reduce the efficacy of selection because an allele that increases/decreases the mutation rate can recombine away from deleterious mutations it generates elsewhere in the genome (Lynch, 2011).

Despite these multiple, interacting factors, previous work has also provided compelling evidence that changing the mutation spectrum alone can drive both the emergence and decay of mutators (Couce et al., 2017, 2013; Couce and Tenaillon, 2019). Here, we provide a mechanistic and general explanation of why, and under what conditions, these effects might occur. In particular, we demonstrate that when a spectrum change either reduces or reverses an existing bias, that is, a bias that has persisted over a period of adaptation, that spectrum change will both increase the beneficial fraction of the DFE and increase the mean selective effect of beneficial mutations.

Our analytical approach, while necessarily simplified, demonstrates mathematically that after a period of adaptation, bias reductions or reversals are expected to increase the beneficial fraction of the DFE, and that this effect will be stronger with larger magnitude reversals. This prediction is confirmed in simulations that relax several simplifying assumptions, treating full populations under both clonal interference and epistasis.

Despite these robust changes to the DFE when the mutation bias is reduced or reversed, a surprising result of our study is that in the absence of changes to mutation rate, the invasion probability of bias-shifted strains is only modestly increased, even under very strong bias reversals (fig. 4, *y*-intercepts). This is because both increases in the beneficial fraction of the DFE and increases in the positive selective effect are of the order of several fold, while mutation rate increases may be 50- or 100-fold or more (Denamur and Matic, 2006). For example, a bias shift that doubles the beneficial fraction (*G* = 2) and increases the positive selective effect by 50% (*H* = 1.5) alters the DFE substantially (fig. 3), but would confer only a three-fold advantage (*GH* = 3) over the wildtype in generating beneficial fixations.

In contrast, as observed for mutations with increased beneficial effect (Couce et al., 2013), bias shifts can dramatically alter the fate of mutator strains (fig. 4). When a shift in the bias is coupled with an increase in mutation rate, the invasion probability of the strain can be vastly increased, including cases in which a disfavoured increase in the mutation rate becomes strongly favoured if coupled with a bias reduction or reversal. This is due to three effects: (1) the expected value of the selection coefficient for beneficial mutations is increased by the bias shift; (2) the fraction of mutations that are beneficial is increased; and (3) as the beneficial fraction is increased, the fraction of mutations that are deleterious is concomitantly reduced. Of these three effects, in our simulated landscapes the increase in beneficial fraction had the greatest magnitude, while the reduction in deleterious load was modest. Further work on empirical fitness landscapes that quantify the DFE for bias-shifted strains (Couce et al., 2013; Sane et al., 2021) is necessary to determine which effects might dominate in natural settings.

We also note that after a change in mutation rate, load accrues gradually, thus mutator strains in our invasion tests may invade before the equilibrium load is realized. This extends the range of mutation rates over which the mutator strain invades beyond the theoretical prediction, which assumed equilibrium load. Since load would also accrue gradually in natural settings after a loss of DNA-repair function, our theoretical prediction is conservative, underestimating the success of bias-shifted mutators. More accurate predictions could include the dynamics of mutational load after a sudden change in mutation rate.

The effects we describe are relevant to asexual evolution, in which changes to mutation rate and/or bias remain linked to the beneficial mutations they generate for sufficient time to reach fixation. Previous work suggests that for mutation rate modifiers, modest levels of horizontal gene transfer or recombination substantially reduce the mutator advantage (Johnson, 1999; Tenaillon et al., 2000). We note that DNA repair enzymes in bacteria are frequently gained and lost, resulting in well-studied changes in mutation rate (Denamur and Matic, 2006; Sane et al., 2021). Changes in mutation spectrum with these gains and losses can also be extreme: a Ti fraction of less than 5% is obtained with the knock-out of the enzyme mutT (Foster et al., 2015), while a Ti fraction of greater than 95% occurs with the loss of mutL (Lee et al., 2012). Invasion simulations that test bias shifts and mutation rates estimated in specific bacterial knock-out strains are thus another clear avenue for future work.

## Acknowledgments

This work was supported by the Natural Sciences and Engineering Research Council of Canada grant RGPIN-2019-06294 and by the National Institute of General Medical Sciences of the National Institutes of Health through grant R01GM127348. We thank Deepa Agashe for helpful discussions.

## Statement of Authorship

MT, DC, RNG and LMW designed the study. LMW, MT and SV wrote the simulation code. MT and SV generated simulation data; MT, DC, RNG and LMW analysed and interpreted simulation data. MT and LMW derived the theory and drafted the manuscript. All authors edited and finalized the manuscript.

## Data and Code Accessibility

Code and data can be accessed at doi:10.5061/dryad.qz612jmk2

