## Supplementary material for "Shifts in mutation bias promote mutators by altering the distribution of fitness effects": Supplemetary Material

### Online Supplement: Shifts in mutation bias promote mutators by altering the distribution of fitness effects, *The American Naturalist*

Marwa Tuffaha[1],
Saranya Varakunan[1],
David Castellano[2],
Ryan N. Gutenkunst[2]
and
Lindi M. Wahl[1,*]

1. Western University, London, Ontario N6A 5B7, Canada;
2. Department of Molecular and Cellular Biology, University of Arizona, Tucson, Arizona 85721, USA.

### Figure S1

Results are shown as for figure 4 in the main text, but here the invasion probability of a single mutant is tested (rather than the invasion probability of a mutant strain at an initial population frequency of 5%).

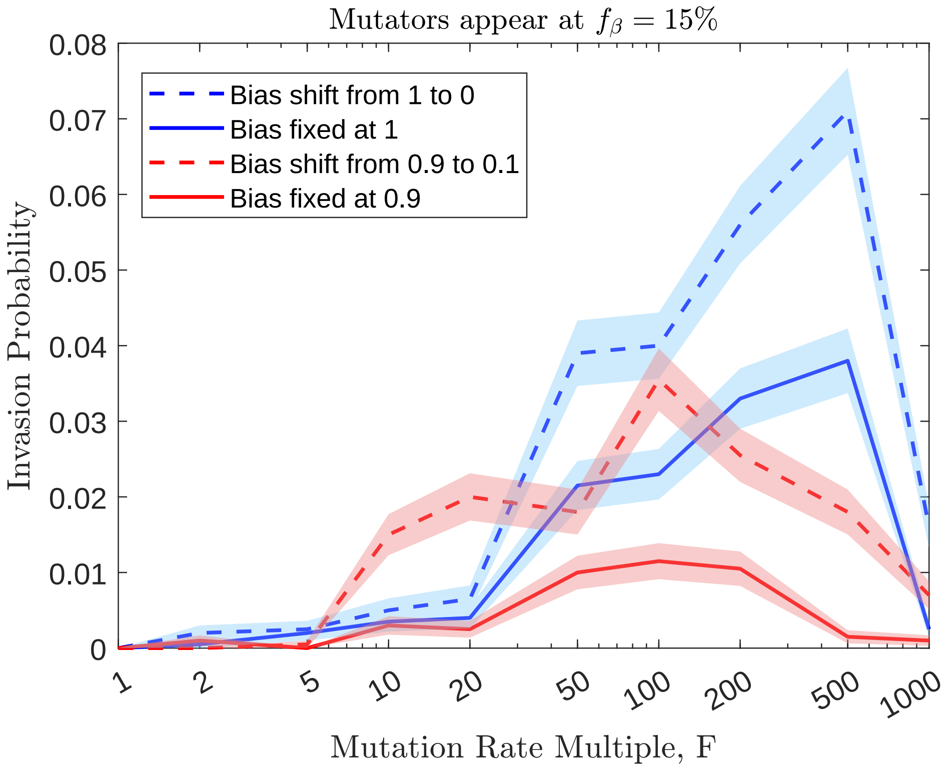


Figure S1: Invasion probabilities for strains with both bias shifts and increased mutation rates (dashed lines) exceed invasion probabilities for strains that increase the mutation rate only (solid lines). Simulation results are shown in which each invading strain is initiated with only one randomly chosen individual from the population and has an $F$-fold increase in the mutation rate ($x$-axis); we compare no change in bias (solid lines), with a full change in bias ($\beta = 1 \rightarrow \beta' = 0$, in blue), or a strong change in bias ($\beta = 0.9 \rightarrow \beta' = 0.1$, in red). Invasion tests occur for populations evolving on an epistatic landscape ($K = 1$) at an adaptive potential of $f_\beta = 15\%$. Results of 2000 replicates are shown; shaded regions indicate $\pm$ one standard deviation

### Figure S2

Changes in deleterious effect size, for parameter values illustrated in figure 3 in the main text ($\beta = 0.7$, $K = 2$).

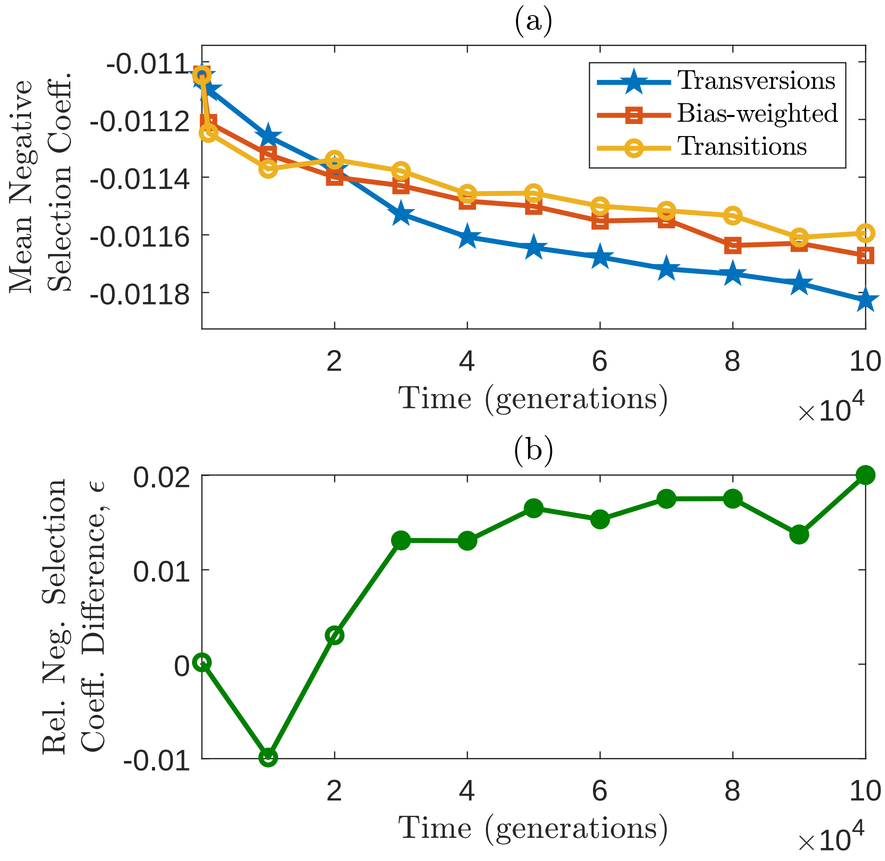


Figure S2: Mean deleterious effect sizes increase in magnitude during the evolution of a transition-biased population. (a) The mean negative selection coefficient of all deleterious transitions (the over-sampled class, yellow circles) decreases over time, falling less rapidly than the analogous value in deleterious transversions (blue stars), except for the first few thousand generations, where noise prevails. The overall mean negative effect size of the bias-weighted DFE (red squares) falls between these two extremes (closer to transitions as expected since the transition frequency $\beta = 0.7$). (b) As a result of their different decay rates, the relative difference in the mean negative selection coefficient, $(s_{\mathrm{Tv}} - s_{\mathrm{Ti}})/s_{\mathrm{Ti}}$ increases over time, but the effect is modest (less than 2% difference).; filled circles indicate that the difference $s_{\mathrm{Tv}} - s_{\mathrm{Ti}}$ is significantly different from zero at that time (t test, $p < .05$). Means of 300 simulation replicates are shown for $\beta = 0.7$ and epistasis degree $K = 2$; DFEs are computed for the most common genotype in the population at each time point. Error bars in top panel are almost equal to symbol heights at most and are omitted.

### Figure S3

Results are shown as for figure 3 in the main text, but without epistasis ($K = 0$).

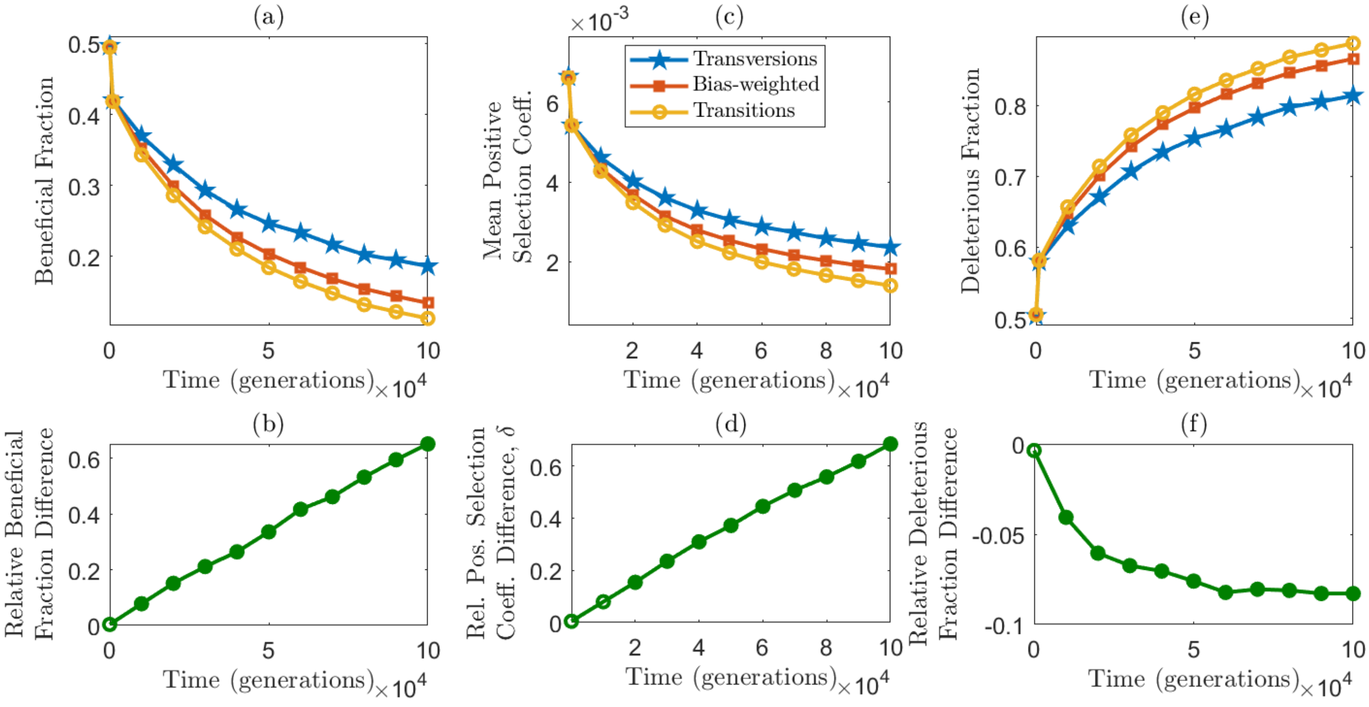


Figure S3: Beneficial fractions and mean beneficial effect sizes decline during the evolution of a transition-biased population, while deleterious fractions increase. (a) The fraction of beneficial transitions (the over-sampled class, yellow circles) decreases over time, falling more rapidly than the fraction of beneficial transversions (blue stars). The overall beneficial fraction of the bias-weighted DFE ($f_\beta$, red squares) falls between these two extremes (closer to $f_{\text{Ti}}$ as expected since the transition frequency $\beta = 0.7$). (b) As a result of their different decay rates, the relative difference in the beneficial fraction, $(f_{\text{Tv}} - f_{\text{Ti}})/f_{\text{Ti}}$ increases over time; filled circles indicate that the difference $f_{\text{Tv}} - f_{\text{Ti}}$ is significantly different from zero at that time (t test, $p < .05$). Panels (c) and (d) show analogous results for the mean positive selection coefficients of transitions and transversions. (e) As the beneficial fraction decreases, the deleterious fraction increases. The fraction of deleterious transitions (yellow circles) increases more rapidly than the fraction of deleterious transversions (blue stars). (f) The relative difference in the deleterious fraction increases in magnitude over time, but the effect is modest (less than 8.5% difference). In all cases, means of 300 simulation replicates are shown for $\beta = 0.7$ and no epistasis ($K = 0$); DFEs are computed for the most common genotype in the population at each time point. Error bars in top panels are smaller than symbol heights and are omitted.

### Figure S4

Results are shown as for figure 3 in the main text, but with epistasis parameter $K = 1$.

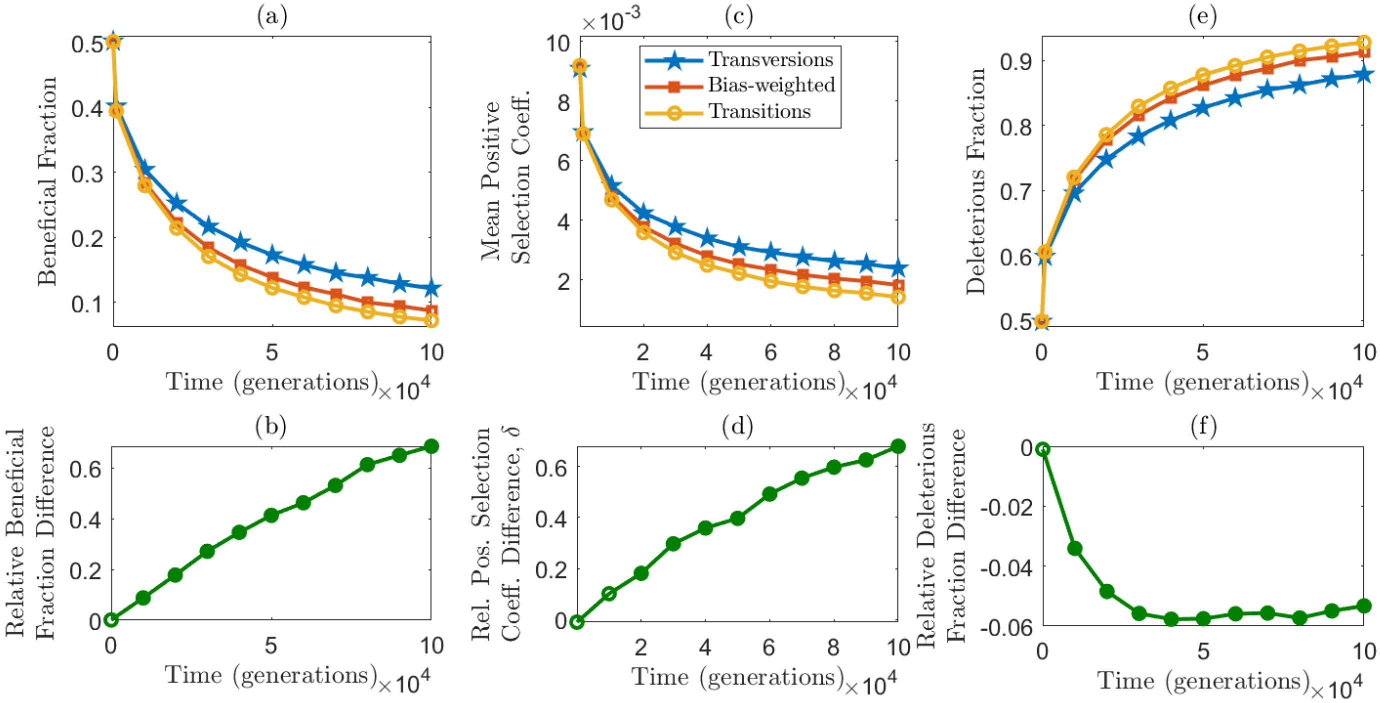


Figure S4: Beneficial fractions and mean beneficial effect sizes decline during the evolution of a transition-biased population, while deleterious fractions increase. (a) The fraction of beneficial transitions (the over-sampled class, yellow circles) decreases over time, falling more rapidly than the fraction of beneficial transversions (blue stars). The overall beneficial fraction of the bias-weighted DFE ($f_\beta$, red squares) falls between these two extremes (closer to $f_{\mathrm{Ti}}$ as expected since the transition frequency $\beta = 0.7$). (b) As a result of their different decay rates, the relative difference in the beneficial fraction, $(f_{\mathrm{Tv}} - f_{\mathrm{Ti}})/f_{\mathrm{Ti}}$ increases over time; filled circles indicate that the difference $f_{\mathrm{Tv}} - f_{\mathrm{Ti}}$ is significantly different from zero at that time (t test, $p < .05$). Panels (c) and (d) show analogous results for the mean positive selection coefficients of transitions and transversions. (e) As the beneficial fraction decreases, the deleterious fraction increases. The fraction of deleterious transitions (yellow circles) increases more rapidly than the fraction of deleterious transversions (blue stars). (f) The relative difference in the deleterious fraction increases in magnitude over time, but the effect is modest (less than 6% difference). In all cases, means of 300 simulation replicates are shown for $\beta = 0.7$ and epistasis degree $K = 1$; DFEs are computed for the most common genotype in the population at each time point. Error bars in top panels are smaller than symbol heights and are omitted.

### Figure S5

Results are shown as for figure 3 in the main text, but with transition frequency $\beta = 0.5$. ($K = 2$).

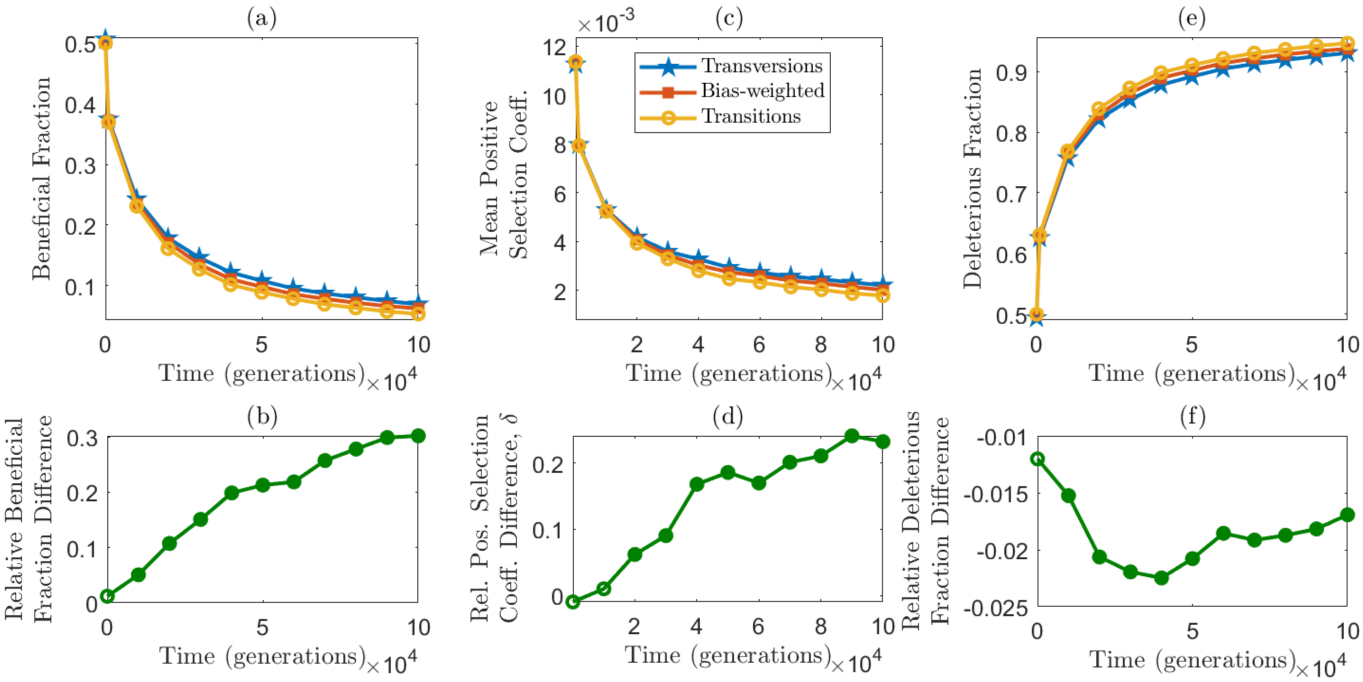


Figure S5: Beneficial fractions and mean beneficial effect sizes decline during the evolution of a slightly transition-biased population, while deleterious fractions increase. (a) The fraction of beneficial transitions (the over-sampled class, yellow circles) decreases over time, falling more rapidly than the fraction of beneficial transversions (blue stars). The overall beneficial fraction of the bias-weighted DFE ($f_\beta$, red squares) falls between these two extremes. (b) As a result of their different decay rates, the relative difference in the beneficial fraction, $(f_{\mathrm{Tv}} - f_{\mathrm{Ti}})/f_{\mathrm{Ti}}$ increases over time; filled circles indicate that the difference $f_{\mathrm{Tv}} - f_{\mathrm{Ti}}$ is significantly different from zero at that time (t test, $p < .05$). Panels (c) and (d) show analogous results for the mean positive selection coefficients of transitions and transversions. (e) As the beneficial fraction decreases, the deleterious fraction increases. The fraction of deleterious transitions (yellow circles) increases more rapidly than the fraction of deleterious transversions (blue stars). (f) The relative difference in the deleterious fraction tends to increase in magnitude over time at the beginning, but the effect is modest (less than 2.5% difference). In all cases, means of 300 simulation replicates are shown for $\beta = 0.5$ and epistasis degree $K = 2$; DFEs are computed for the most common genotype in the population at each time point. Error bars in top panels are smaller than symbol heights and are omitted.

### Figure S6

Results are shown as for figure 3 in the main text, but with transition frequency $\beta = 0.1$. Here $K = 2$ as in figure 3.

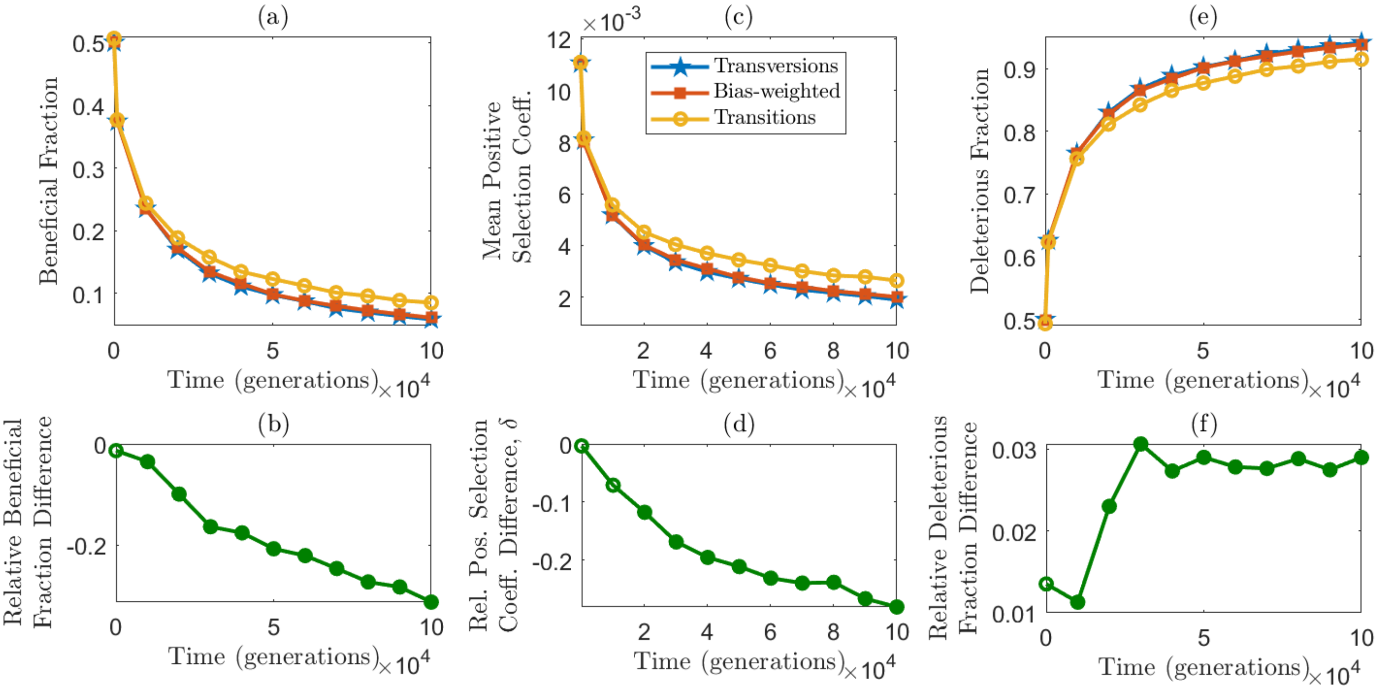


Figure S6: Beneficial fractions and mean beneficial effect sizes decline during the evolution of a transversion-biased population, while deleterious fractions increase. (a) The fraction of beneficial transversions (the over-sampled class, blue stars) decreases over time, falling more rapidly than the fraction of beneficial transitions (yellow circles). The overall beneficial fraction of the bias-weighted DFE ($f_\beta$, red squares) falls between these two extremes (closer to $f_{\text{Tv}}$ as expected due to the transition frequency $\beta = 0.1$). (b) As a result of their different decay rates, the relative difference in the beneficial fraction, $(f_{\text{Tv}} - f_{\text{Ti}})/f_{\text{Ti}}$ decreases over time; filled circles indicate that the difference $f_{\text{Tv}} - f_{\text{Ti}}$ is significantly different from zero at that time (t test, $p < .05$). Panels (c) and (d) show analogous results for the mean positive selection coefficients of transitions and transversions. (e) As the beneficial fraction decreases, the deleterious fraction increases. The fraction of deleterious transversions (blue stars) increases more rapidly than the fraction of deleterious transitions (yellow circles). (f) The relative difference in the deleterious fraction tends to increase in magnitude over time, but the effect is modest (less than 3.5% difference). In all cases, means of 300 simulation replicates are shown for $\beta = 0.1$ and epistasis degree $K = 2$; DFEs are computed for the most common genotype in the population at each time point. Error bars in top panels are smaller than symbol heights and are omitted.

### Figure S7

Results are shown as for figure 5 in the main text, but for a mutator with a 10-fold increase in mutation rate ($F = 10$).

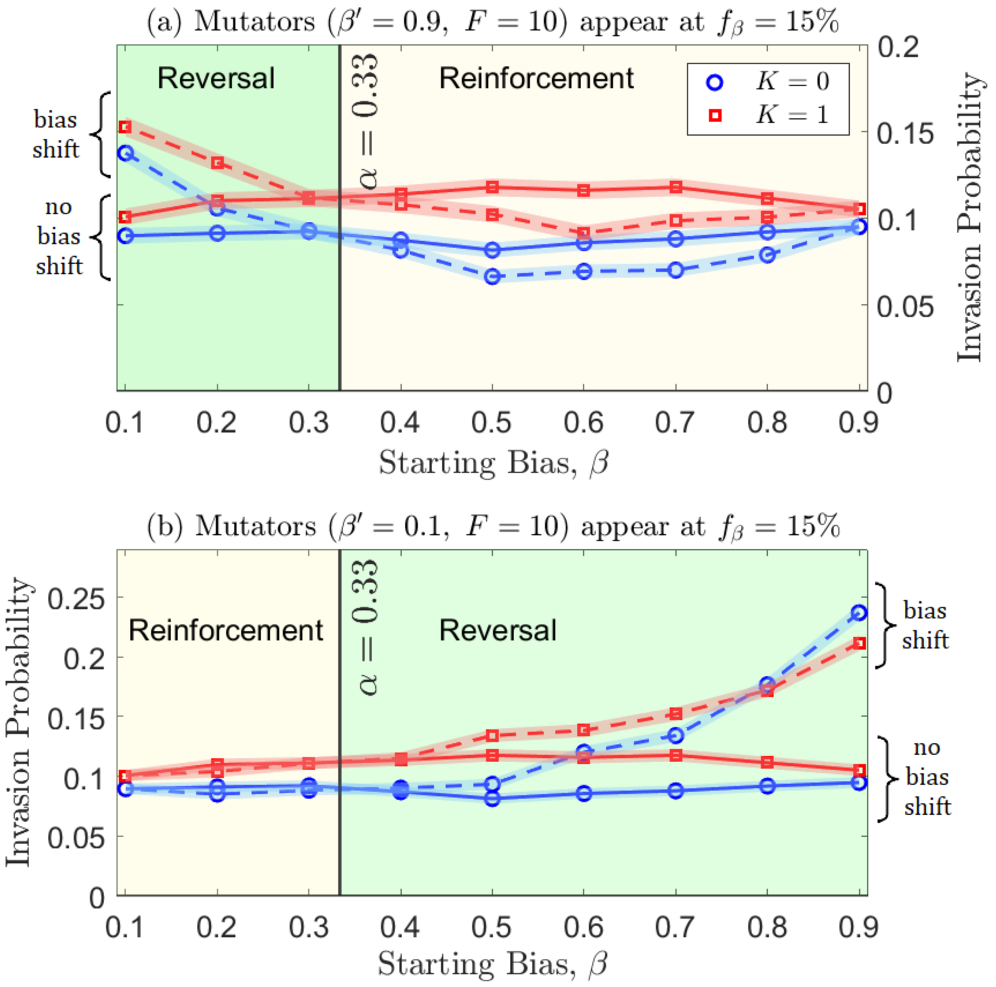


Figure S7: Invasion probabilities of mutators increase when the bias is reversed. Populations evolved with a fixed starting bias ($x$-axis) until the beneficial fraction of the most common genotype reached 15%. An invasion test was then simulated for a strain with a 10-fold increase in mutation rate. Two cases are compared: the mutators either have the same bias as the initial population (solid lines), or have a bias shift to (a) $\beta' = 0.9$ or (b) $\beta' = 0.1$ (dashed lines). The vertical line at $\alpha = 1/3$ represents the unbiased transition frequency. Panel (a) shows that for a starting bias less than $\alpha$, a shift to $\beta' = 0.9$ reverses the bias and thus increases the invasion probability (dashed lines above solid lines); the same bias shift tends to reduce the invasion probability (dashed lines below solid lines) if the starting bias is greater than $\alpha$. The effects are reversed when the bias shifts to $\beta' = 0.1$, as shown in panel (b). Results of 4000 replicates are shown; shaded regions indicate $\pm$ one standard deviation.

### Figure S8

Results are shown as for figure 5 in the main text, but here invasions are tested at adaptive potential $f_\beta = 30\%$.

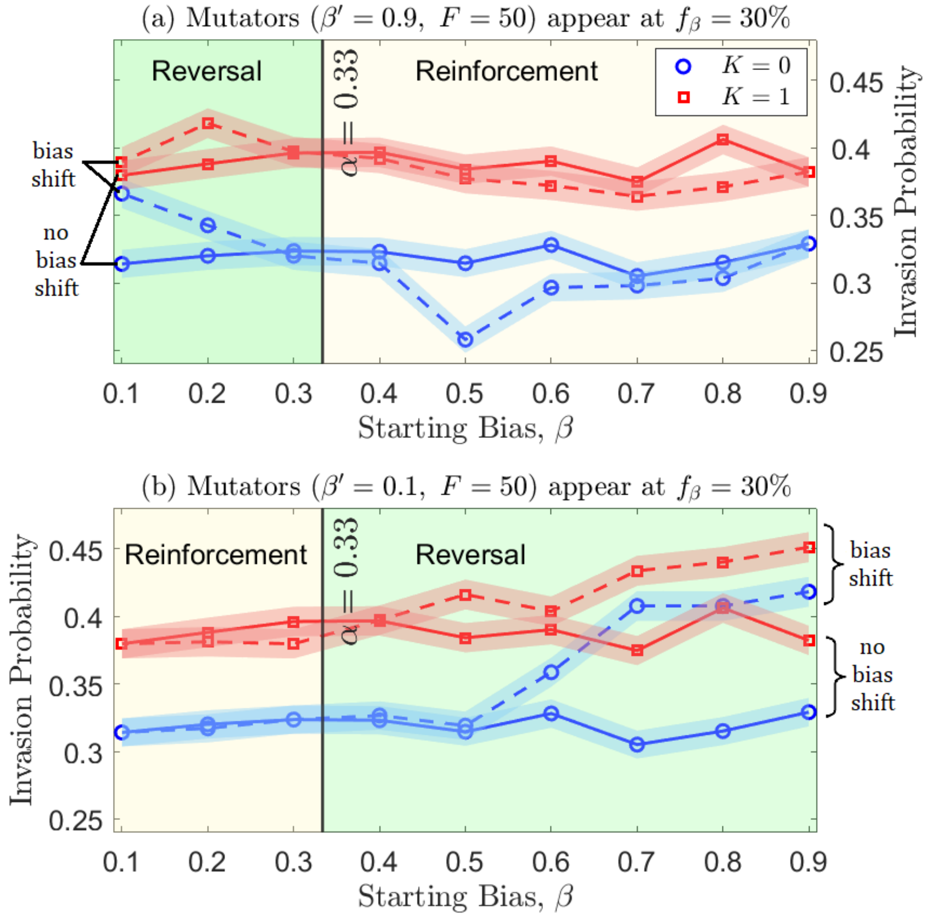


Figure S8: Invasion probabilities of mutators increase when the bias is reversed. Populations evolved with a fixed starting bias ($x$-axis) until the beneficial fraction of the most common genotype reached 30%. An invasion test was then simulated for a strain with a 50-fold increase in mutation rate. Two cases are compared: the mutators either have the same bias as the initial population (solid lines), or have a bias shift to (a) $\beta' = 0.9$ or (b) $\beta' = 0.1$ (dashed lines). The vertical line at $\alpha = 1/3$ represents the unbiased transition frequency. Panel (a) shows that for a starting bias less than $\alpha$, a shift to $\beta' = 0.9$ reverses the bias and thus increases the invasion probability (dashed lines above solid lines); the same bias shift tends to reduce the invasion probability (dashed lines below solid lines) if the starting bias is greater than $\alpha$. The effects are reversed when the bias shifts to $\beta' = 0.1$, as shown in panel (b). Results of 2000 replicates are shown; shaded regions indicate $\pm$ one standard deviation.

### Figure S9

Results are shown as for figure 5 in the main text, but for a mutator with a 200-fold increase in mutation rate ($F = 200$) and at an adaptive potential of $f_\beta = 30\%$.

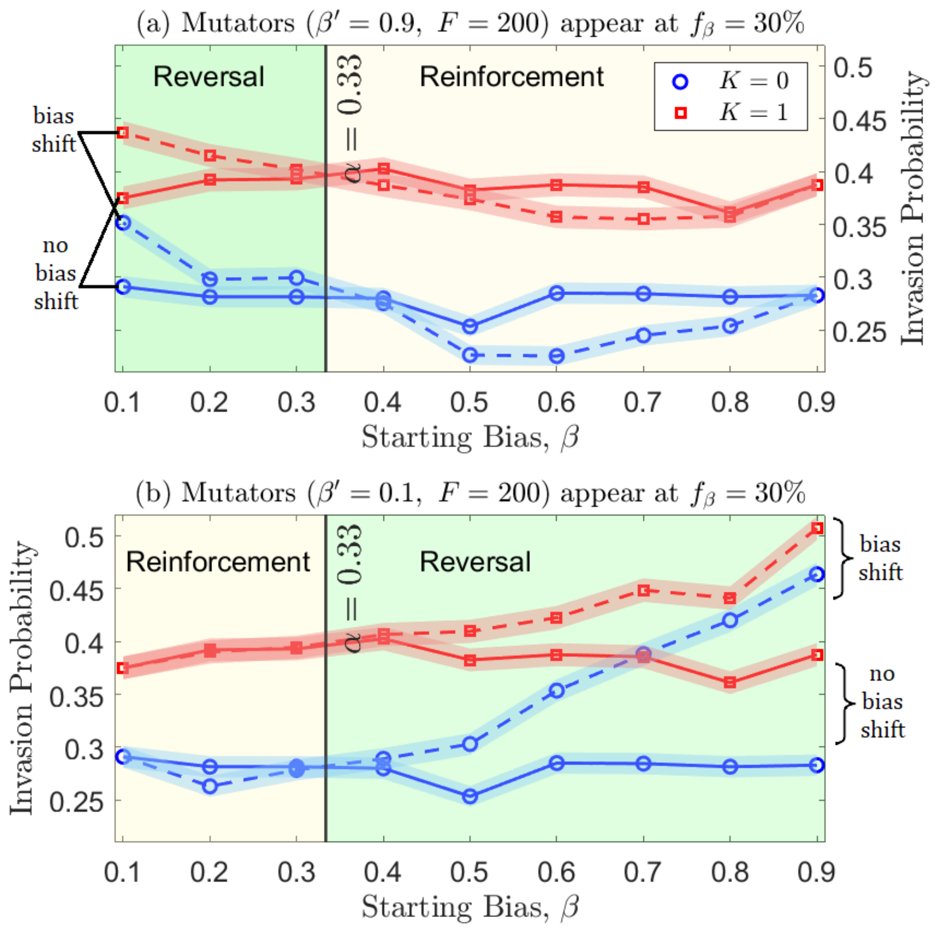


Figure S9: Invasion probabilities of mutators increase when the bias is reversed. Populations evolved with a fixed starting bias ($x$-axis) until the beneficial fraction of the most common genotype reached 30%. An invasion test was then simulated for a strain with a 200-fold increase in mutation rate. Two cases are compared: the mutators either have the same bias as the initial population (solid lines), or have a bias shift to (a) $\beta' = 0.9$ or (b) $\beta' = 0.1$ (dashed lines). The vertical line at $\alpha = 1/3$ represents the unbiased transition frequency. Panel (a) shows that for a starting bias less than $\alpha$, a shift to $\beta' = 0.9$ reverses the bias and thus increases the invasion probability (dashed lines above solid lines); the same bias shift tends to reduce the invasion probability (dashed lines below solid lines) if the starting bias is greater than $\alpha$. The effects are reversed when the bias shifts to $\beta' = 0.1$, as shown in panel (b). Results of 2000 replicates are shown; shaded regions indicate $\pm$ one standard deviation.
